# Cellular senescence in malignant cells promotes tumor progression in mouse and patient Glioblastoma

**DOI:** 10.1101/2022.05.18.492465

**Authors:** Rana Salam, Alexa Saliou, Franck Bielle, Mathilde Bertrand, Christophe Antoniewski, Catherine Carpentier, Agusti Alentorn, Laurent Capelle, Marc Sanson, Emmanuelle Huillard, Léa Bellenger, Justine Guégan, Isabelle Le Roux

## Abstract

Glioblastoma (GBM) is the most common primary malignant brain tumor in adults, yet it remains refractory to systemic therapy. Elimination of senescent cells has emerged as a promising new treatment approach against cancer. Here, we investigated the contribution of senescent cells to GBM progression. Senescent cells were identified in patient and mouse GBMs. Partial removal of p16^Ink4a^-expressing malignant senescent cells, which make up less than 7 % of the tumor, modified the tumor ecosystem and improved the survival of GBM-bearing mice. By combining single cell and bulk RNA sequencing, immunohistochemistry and genetic knockdowns, we identified the NRF2 transcription factor as a determinant of the senescent phenotype. Remarkably, our mouse senescent transcriptional signature and underlying mechanisms of senescence are conserved in patient GBMs, in whom higher senescence scores correlate with shorter survival times. These findings suggest that senolytic drug therapy may be a beneficial adjuvant therapy for patients with GBM.

## Introduction

Diffuse gliomas are the most common primary malignant brain tumors in adults^1^. Glioblastoma (GBM; IDH-wild type glioma, grade 4) is the most aggressive glioma and despite intensive conventional therapy which includes surgery, radiation and both concurrent and adjuvant temozolomide (TMZ) chemotherapy, GBM remains treatment resistant and disease progression is fatal, with a median survival below 15 months^2^. Distinct factors may account for current treatment failures, including tumor invasiveness, an immunosuppressive microenvironment and intra-tumoral heterogeneity. Novel approaches are therefore required to find effective therapeutic strategies.

Cellular senescence is a permanent cell cycle arrest mediated by p53/p21^CIP1^ and/or p16^INK4A^/Rb pathways and is defined by a combination of features including a senescence-associated secretory phenotype (SASP), anti-apoptotic program, and increased lysosomal content, the latter allowing histochemical detection of senescence associated-β-galactosidase activity (SA-β-gal)^3^. In cancer, cellular senescence is triggered by multiple stresses such as DNA damage, oncogene activation, therapeutic agents or elevated reactive oxygen species (ROS). SASP is defined by the secretion of a plethora of factors including cytokines, chemokines, growth factors, extracellular matrix (ECM) components and proteases, which together can stimulate angiogenesis, modulate the composition of the ECM and promote an epithelial-to-mesenchymal transition^4,5^. Depending on the context, senescence exerts two opposite effects during tumorigenesis^6^. In some contexts, senescent cells prevent the proliferation of pre-malignant cancer cells, as SASP factors stimulate the immune clearance of oncogene or therapy-induced senescent tumor cells^7,8,9^. Conversely, in persistently senescent cells, the SASP can either directly induce tumor growth^10^ or contribute to immune suppression, thus allowing tumor progression^11,12^. Many studies have assessed the function of senescence in developing tissues and age-related diseases by the *in vivo* removal of senescent cells, either using chemical or genetic senolytics^13,14,15,16,17^. A common genetic approach employs *p16^Ink4a^* regulatory sequences to drive the inducible expression of INK-ATTAC or p16-3MR, which selectively eliminate senescent cells expressing high levels of *p16^Ink4a^*, leading to apoptosis^16,17^. This senolytic strategy efficiently reduces the adverse effects of therapy-induced senescent cells in a mouse breast cancer model^18^.

A few *in vivo* studies have begun to examine the role of cellular senescence in gliomas. Using mouse patient-derived xenograft (PDX) models, it was shown that IL6, a universal SASP component as well as a cytokine express by immune cells, promotes growth of patient glioma stem cells (GSCs) and contributes to glioma malignancy^19^. Conversely, loss of PTEN-PRMT5 signaling induces senescent GSCs to slow down tumorigenesis^20^. Furthermore, loss of one allele of p53 reduces H-RasV12 oncogene-induced senescence in an orthotopic GBM model, as evidenced by reduced SA-β-gal staining, and decreases mouse survival time^21^. Finally, a recent study revealed the dual effect of therapy-induced senescence (TIS) following BMI1 inhibitor treatment of diffuse intrinsic pontine glioma tumor (a pediatric high-grade glioma), which initially attenuates tumor cell self-renewal and growth, but later leads to SASP-mediated tumor recurrence^22^. This study confirmed the detrimental function of persistent senescent cells in glial tumors, and suggested that senescent cells could represent an actionable target to mitigate the process of gliomagenesis^6^.

Recent single-cell RNA sequencing studies classified the intra-tumoral heterogeneity of malignant GBM cells^23,24,25,26,27^, which can be subdivided into four main cellular plastic states: oligodendrocyte progenitor cell-like (OPC-like), neural progenitor cell-like (NPC-like), astrocyte-like (AC-like) and mesenchymal-like (MES-like) states^23^. The relative abundance of these cellular states within the tumor defines three GBM transcriptomic subtypes, with proneural (PN-GBM) and classical (CL-GBM) GBMs associated with neurodevelopmental programs and mesenchymal GBM (MES-GBM) associated with injury response programs^24,25,26,27,28,29^. OPC-like and NPC-like states are enriched in PN-GBM whereas AC-like and MES-like states are enriched in CL-GBM and MES-GBM, respectively^23^. Notably, stemness programs are heterogeneous even within a single tumor and PN- and MES-GSCs could contribute to the genetic heterogeneity observed in patient GBM^24,26,30^. Each transcriptional GBM subtype is associated with distinct molecular alterations and patient outcomes. MES-GBM is correlated with enhanced activation of anti-inflammatory (or tumor-promoting) macrophages^29,31,32,33^. Mutations in *NF1, TP53, PTEN* genes and increased NF-κB signaling are prevalent in this GBM subtype^28^. Interestingly, *PTEN* loss induces cellular senescence and activates NF-κB signaling, which initiates and maintains the SASP^34,35,36^. Together these findings support the idea that cellular senescence could contribute to the intra-tumoral heterogeneity of GBM.

In this study, we investigated whether cellular senescence participates in GBM tumor progression using patient-resected GBM tissues and a mouse GBM model^37^. We identified senescent cells in patient and mouse GBMs. Partial removal of senescent cells expressing high levels of p16^Ink4a^ using a ganciclovir-inducible p16-3MR transgenic line^17^ improved the survival of GBM-bearing mice. To identify the cells expressing high levels of p16^Ink4a^, and to characterize the action of these cells on the tumor ecosystem, we combined single cell and bulk RNA sequencing (RNAseq) analysis at early and late timepoints after the senolytic treatment. This approach led to the identification of the NRF2 transcription factor and its selected targets as a signal triggering the pro-tumoral activity of p16^Ink4a^ expressing senescent cells. Using these data, we defined an unbiased senescence signature that we successfully used to interrogate GBM patient data sets, revealing that higher senescence scores correlated with shorter survival times.

## Results

### Identification of senescent cells in patient and mouse gliomas

We first searched for senescent cells in 28 freshly resected diffuse gliomas from patients by performing SA-β-gal staining coupled with immunohistochemistry (IHC) (14 GBMs, 5 astrocytomas, 9 oligodendrogliomas; Supplementary Fig. 1a). Senescent cells were identified as SA-β-gal positive (SA-β-gal+) and negative for the cell cycle marker Ki67 (Ki67-; Fig.1a). Depending on the molecular alterations found in gliomas, some SA-β-gal+ cells expressed the cell cycle inhibitor p16^INK4A^, while in gliomas harboring a mutation of p53, senescent malignant cells expressed high levels of mutant p53 (Fig.1a and Supplementary 1a). To identify senescent cells, we used cell type-specific markers. Some SA-β-gal+ cells co-expressed GFAP, which could either represent parenchymal astrocytes or tumor cells, OLIG2, an OPC marker, or IBA1, a microglia/macrophage marker (Fig. 1a). To establish a quantitative measure of senescent cell burden, we quantified the percentage of the tumor area containing SA-β-gal+ cells, and used these measures to stratify tumors into 3 senescent categories: (1) >1% (n=10/28) but below 7%; (2) ≤1-0.1%> (n=13/28) and (3) ≤0.1% (n=5/28) senescent cells (Supplementary Fig. 1a and b). Notably, no diffuse glioma types were associated with a particular senescent category.

**Figure 1.**
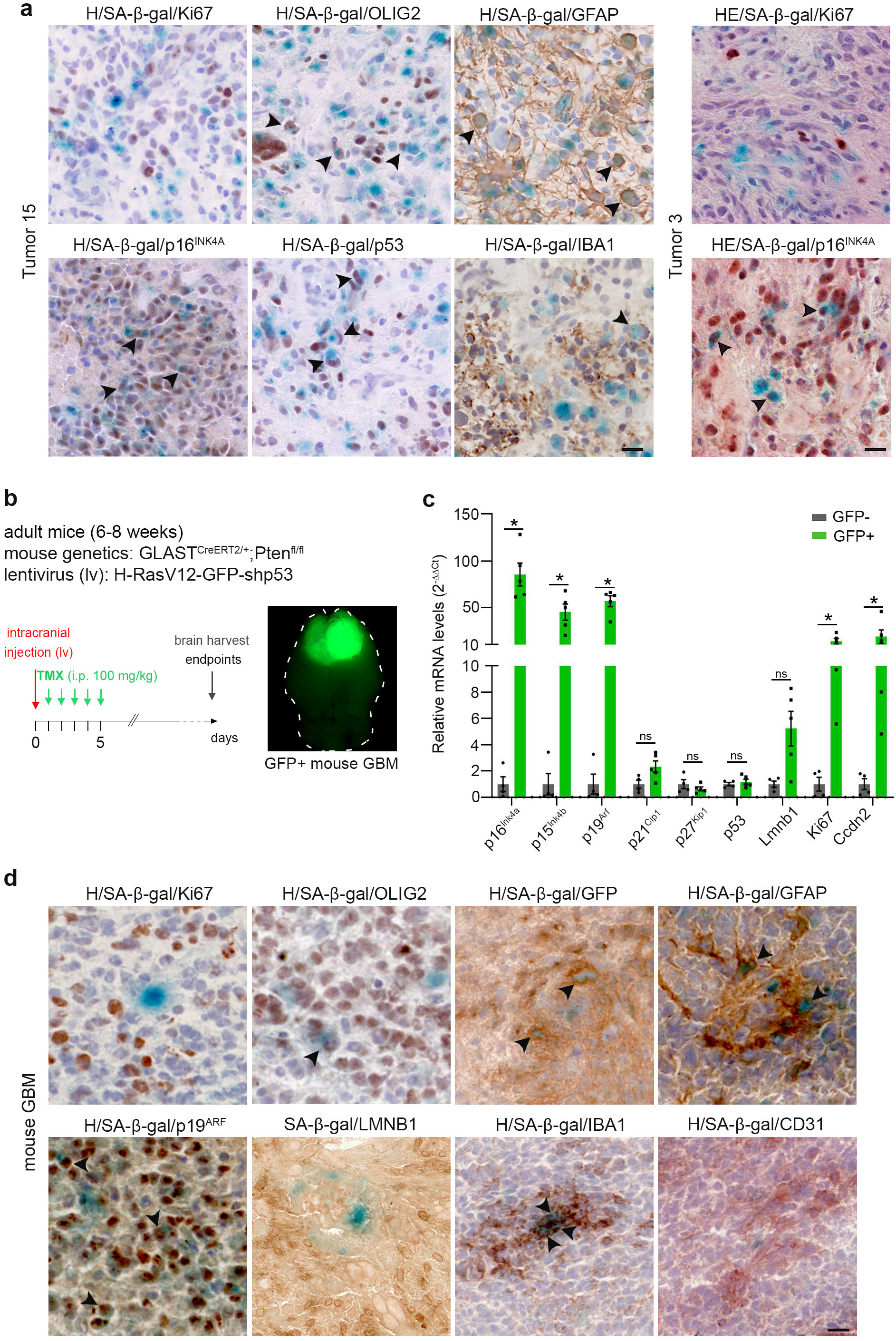
Identification of senescent cells in patient and mouse gliomas. (**a**) Representative SA-β-gal staining (blue) coupled with immunohistochemistry (IHC, brown) on two non-fixed patient GBM cryosections samples (Ki67 and GFAP: n=16; p16^INK4A^: n=12; IBA1: n=10; p53: n=7 and OLIG2: n=6 patient GBMs). (**b**) Left: genetics of the mouse mesenchymal GBM model (mouse and injected lentivirus (lv)). The time line represents the induction of the tumorigenesis with tamoxifen intraperitoneal (i.p.) injections (TMX, 100mg/kg/day for 5 days). Brains are harvested when mice reach end points. Right: representative stereomicroscopic image of a mouse brain with a GFP+ GBM. (**c**) Relative transcript levels shown as ratios of normalized values of mouse GBM (GFP+, n=4) over surrounding parenchyma (GFP-, n=4). Data are represented as the mean ± SEM. Statistical significance was determined by Wilcoxon-Mann-Whitney test (*, p<0.05; ns, not significant). (**d**) Representative SA-β-gal staining (blue) coupled with IHC (brown) on mouse GBM cryosections. (Ki67, p19, IBA1 and GFP: n=8; GFAP: n=6; OLIG2 and CD31: n=5; LMNB1: n=4 independent mouse GBMs). Arrow heads in **a**, **d** point to double positive cells. Scale bars, **a** and **d**: 20 μm. H: hematoxylin; HE: hematoxylin and eosin; i.p.: intraperitoneal; TMX: tamoxifen.

We next investigated whether specific molecular alterations were associated with each of the senescent tumor categories, as defined by SA-β-Gal cell percentages. Homozygous deletion of *CDKN2A*, encoding for p16^INK4A^, is carried by 54% of patient GBMs (cbioportal.org). As p16^INK4A^ is a mediator of senescence, we annotated p16^INK4a^ status of each tumor, as well as examining other common molecular alterations, including p53, PTEN, NF1, and EGFR mutations. Notably, we did not find any association of a specific molecular alteration with a single senescent category (Supplementary Fig. 1a).

Finally, we studied senescence in an immuno-competent GBM mouse model, employing a modified version of a model developed by Friedmann-Morvinski *et al*.^37^. This model recapitulates the molecular alterations identified in MES-GBM: the loss of *Pten* and *p53* and the inactivation of *Nf1* triggered by the ectopic expression of H-RasV12 (Fig. 1b). Six-to-eight week-old Glast^creERT2/+^;Pten^fl/fl^ mice were intracranially injected with a lentivirus encoding H-RasV12-IRES-eGFP and shp53, into the subventricular zone (SVZ). Mice were sacrificed when they reached disease end points, hereafter referred to as late timepoint (Fig. 1b). These tumors displayed a heterogenous histopathology similar to that described in patient GBMs^38^ (Supplementary Fig. 1c). By qPCR analysis, elevated expression levels of *Ink4/ARF* (encoding p16^Ink4a^, p15^Ink4b^, p19^Arf^) and *p21*, both of which encode senescence-mediating proteins, were detected in the tumor (GFP+) cells compared with the surrounding parenchyma (GFP-) cells (Fig. 1c). In contrast, *p53* mRNA levels were similarly low within GFP+ tumor cells and adjacent GFP-cells, in agreement with the presence of shp53 in the lentivirus (Fig. 1c). Notably, p16^Ink4a^ protein could not be examined in mouse tissues due to the lack of a suitable antibody. However, using immunohistochemistry, we identified SA-β-gal+ Ki67-LAMINB1- p19^ARF^+ senescent cells in mouse GBMs (Fig. 1d). These senescent cells were of distinct types, and included either malignant (GFP+) tumor cells, glial cells (GFAP+, OLIG2+), or microglia/macrophage (IBA1+) (Fig.1d). We did not detect any senescent endothelial cells (CD31+; Fig. 1d). In general, the senescent cells were sparsely distributed in the tumor, and mostly located in proliferative areas or adjacent to necrotic regions (Supplementary Fig. 1d).

All together these data reveal that cellular senescence is associated with primary gliomagenesis, including in the mouse GBM model, which recapitulates the histopathology, senescence features and cell identities of patient GBMs. We thus further used this model to address the function of senescent cells during primary gliomagenesis.

### Senescent cells partial removal increases the survival of GBM-bearing mice

We introduced the p16-3MR transgene in the mouse GBM model, which allowed us to selectively remove senescent cells expressing high levels of *p16^Ink4a^* with ganciclovir (GCV) injections^17^. Remarkably, the median survival of GBM-bearing mice harboring p16-3MR that were treated with GCV (p16-3MR+GCV) increased significantly compared with WT mice treated with GCV (WT+GCV) or p16-3MR mice treated with vehicle (p16-3MR+vhc) (Fig. 2a-c). Similarly, the survival of GBM-bearing mice treated with the senolytic drug ABT263 (Navitoclax, an inhibitor of the anti-apoptotic proteins BCL2 and BCL-xL^13^) increased significantly compared with control mice (WT+vhc) (Fig. 2b and d). Together these results strongly suggest that senescent cells act as a pro-tumoral mechanism during primary gliomagenesis.

**Figure 2.**
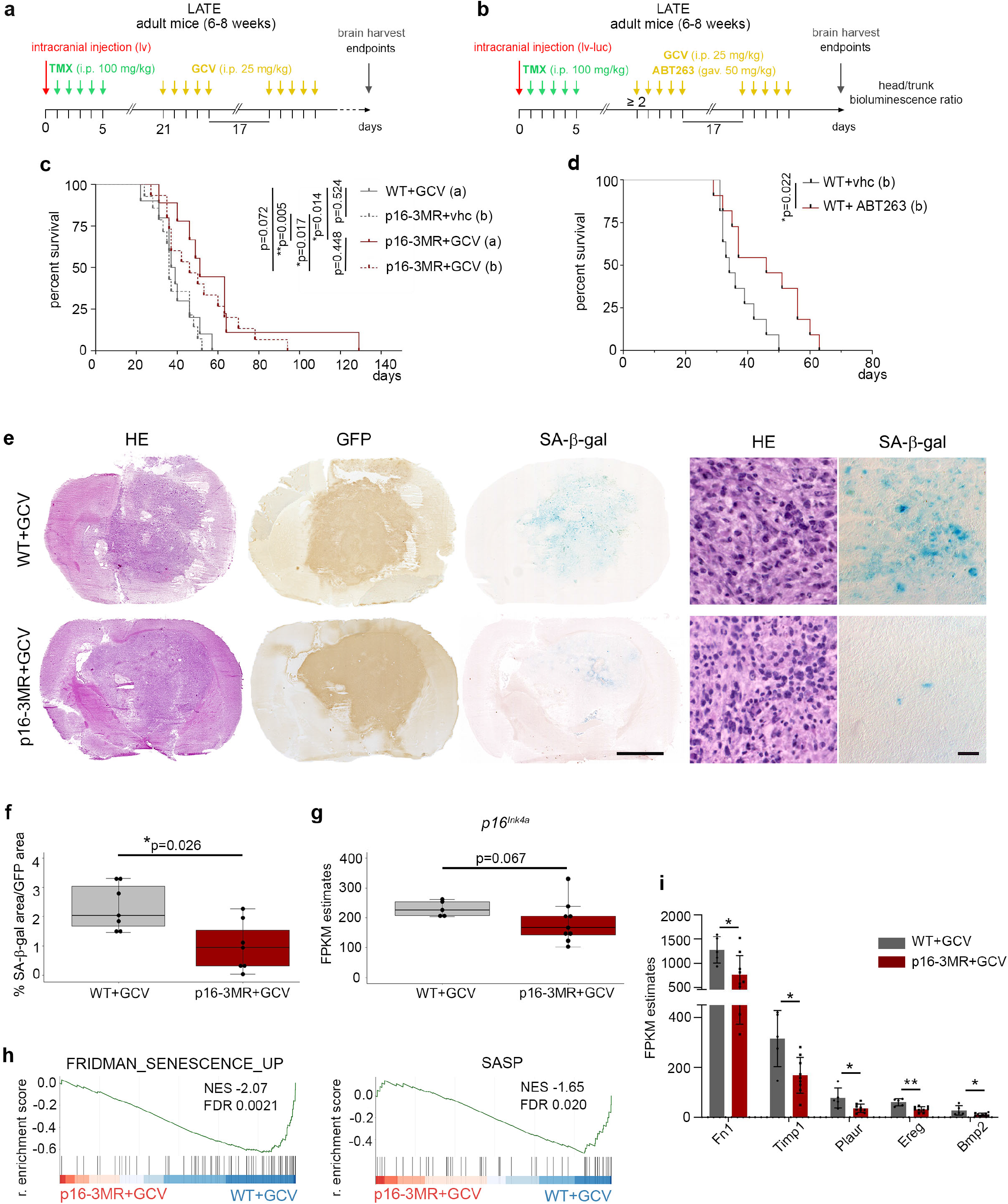
Senescent cells partial removal increases the survival of GBM-bearing mice. (**a**) Timeline of tumorigenesis induction (lv: H-RasV12-GFP-shp53) and removal of p16^Ink4a^ senescent cells with GCV i.p. injections, 21 days post lv injection in the p16-3MR transgenic mouse. (**b**) Timeline of tumorigenesis induction (lv-luc: H-RasV12-GFP-P2A-Luc2-shp53) and removal of senescent cells with GCV i.p. injections in the p16-3MR transgenic mouse or with ABT263 gavage in WT mouse when head to body bioluminescence ratio reached 2. (**c**) Kaplan-Meier survival curves (solid lines) of WT (n=10, median survival 38 days) and p16-3MR (n=9, median survival 51 days) mice treated with GCV as shown in **a**. Kaplan-Meier survival curves (dotted lines) of p16-3MR mice treated with vhc (n=14, median survival 36 days) or GCV (n=15, median survival 46 days) as shown in **b**. (**d**) Kaplan-Meier survival curves of WT mice treated with vhc (n=11, median survival 34 days) or ABT263 (n=11, median survival 46 days) as shown in **b**. (**e**) Representative HE, GFP IHC and SA-β-Gal staining on adjacent mouse GBM cryosections. GFP IHC was used to delineate the tumor area. Right panels represent higher magnifications of the left panels. Scale bars, left panels: 2.5 mm, right panels: 20 μm. (**f**) Quantification of the SA-β-Gal area over the tumor (GFP+) area. (**g**) Relative transcript levels of *p16^Ink4a^*, shown as FPKM estimates extracted from the bulk RNAseq analysis (WT+GCV, n=5; p16-3MR+GCV, n=9). (**h**) GSEA graphs from bulk RNAseq data, representing the enrichment score of two senescence pathways in p16-3MR+GCV GBMs compared with WT+GCV GBMs. The barcode plot indicates the position of the genes in each gene set; red represents positive Pearson’s correlation with p16-3MR+GCV expression, blue with WT+GCV expression. The SASP gene list is a custom list established from Gorgoulis *et al*.^3^ (Supplementary Table 1). (**i**) Relative transcript levels of genes in WT+GCV and p16-3MR+GCV GBMs; FPKM estimates were extracted from bulk RNAseq data. These genes are differentially expressed between the two conditions. **c, d** Statistical significance was determined by Mantel-Cox log-rank test (*, p<0.05, **, p<0.01). **f**, **g**, **i** Data are represented as the mean ± SD and statistical significance was determined by Wilcoxon-Mann-Whitney test (*, p<0.05; **, p<0.01). TMX: tamoxifen; vhc: vehicle; gav.: gavage; GCV: ganciclovir; i.p.: intraperitoneal; lv: lentivirus; lv-luc: lentivirus-luciferase; HE: hematoxylin and eosin; GSEA: gene set enrichment analysis; FDR: false discovery rate; NES: normalized enrichment score; r. enrichment score: running enrichment score.

To confirm the tumor promoting function of senescent cells, we further studied GBM mice carrying the p16-3MR transgene. First, we analyzed whether senescence hallmarks decreased in p16-3MR+GCV GBMs compared with controls at the late timepoint (i.e., disease endpoint). We quantified the percentage of the tumor area (defined by GFP expression) encompassing SA-β-gal cells, and found that it decreased 2.3-fold (from 2.32% to 0.99%) in p16-3MR+GCV tumors compared to WT+GCV GBMs (Fig. 2e and f). On average, about 2% of the tumor area was comprised of SA-β-gal cells in WT+GCV GBMs, which corresponds to senescent category one as we defined using patient gliomas (Supplementary Fig. 1a).

We next performed bulk RNA sequencing (RNAseq) of the tumors with or without senescent cells. In agreement with the inter-tumoral heterogeneity of patient GBMs, heatmaps of the bulk RNAseq data revealed inter-tumoral heterogeneity of mouse GBMs independent of the treatment (Supplementary Fig. 2a and c). Gene Set Enrichment Analysis (GSEA, Supplementary Fig. 2e) of p16-3MR+GCV GBMs compared with WT+GCV GBMs revealed an upregulation of cell cycle components (E2F targets), a downregulation of pathways involved in cancer (Notch signaling, mTORC1 signaling, epithelial-mesenchymal transition, angiogenesis), and a modulation of the immune system (TNFA signaling via NFKB, Interferon responses, Il2-Stat5 signaling). In addition, bulk RNAseq analysis revealed a slight decrease although no significant, in *p16^Ink4a^* transcripts in p16-3MR+GCV GBMs compared with control GBMs (Fig. 2g). Finally, GSEA revealed a significant downregulation of senescence pathways (Fig. 2h, Supplementary Fig. 2f; Supplementary Table 1). SASP genes whose expression was significantly decreased in p16-3MR+GCV compared with WT+GCV GBMs included *Fn1, Plau, Timp1, Ereg*, and *Bmp2*^39,40,41^ (Fig. 2i, Supplementary Fig. 2b and d). These SASP genes encode for growth factors and extracellular matrix components or remodelers.

Collectively our data show that at the late timepoint, when mice were sacrificed due to tumor burden, there was an increased survival of GBM bearing mice associated with the partial removal of p16^Ink4a^ senescent cells, therefore pointing to the tumor promoting action of senescent cells during gliomagenesis.

### Identification of p16^Ink4a Hi^ cells in a subset of malignant cells

To unveil the identity of senescent cells expressing high levels of *p16^Ink4a^* and targeted by the p16-3MR transgene with GCV^17^, we performed droplet-based single cell RNAseq (scRNAseq) on FACs sorted cells from WT and p16-3MR GBM cells collected 7 days after the last GCV injection, hereafter named early timepoint (Fig. 3a and b). At this stage, WT+GCV GBMs (n=2) exhibit increased tumor growth compared with p16-3MR+GCV GBMs (n=2) (Supplementary Fig. 3a-c). Uniform manifold approximation and projection (UMAP) clustering at 0.5 resolution revealed 22 clusters with distinct gene expression signatures in each sample in the two conditions (Fig. 3c and Supplementary Fig.3d). Non-malignant cells were identified based on the expression of the pan-leucocyte marker *CD45* (*Ptprc*), and malignant cells were identified by their expression of the 3’ long terminal repeat (*3’LTR*) of the injected lentivirus and by copy number variations (CNV) (Fig. 3d and Supplementary Fig. 3e). Cells in each of the 22 clusters expressed variable levels of *p16^Ink4a^* (*Cdkn2a*) however, only malignant tumor cells expressed high levels of *p16^Ink4a^*. Hereafter, we refer to p16^Ink4a Hi^ cells as those cells expressing *p16^Ink4a^* at a level ≥ 4 (Fig. 3d). This cut off was chosen as p16^Ink4a Hi^ cells represent 3% (412/13563) of the tumor cells, a percentage that is in agreement with the area of SA-β-Gal staining in the tumors (Supplementary Fig. 1a and Fig. 2f).

**Figure 3.**
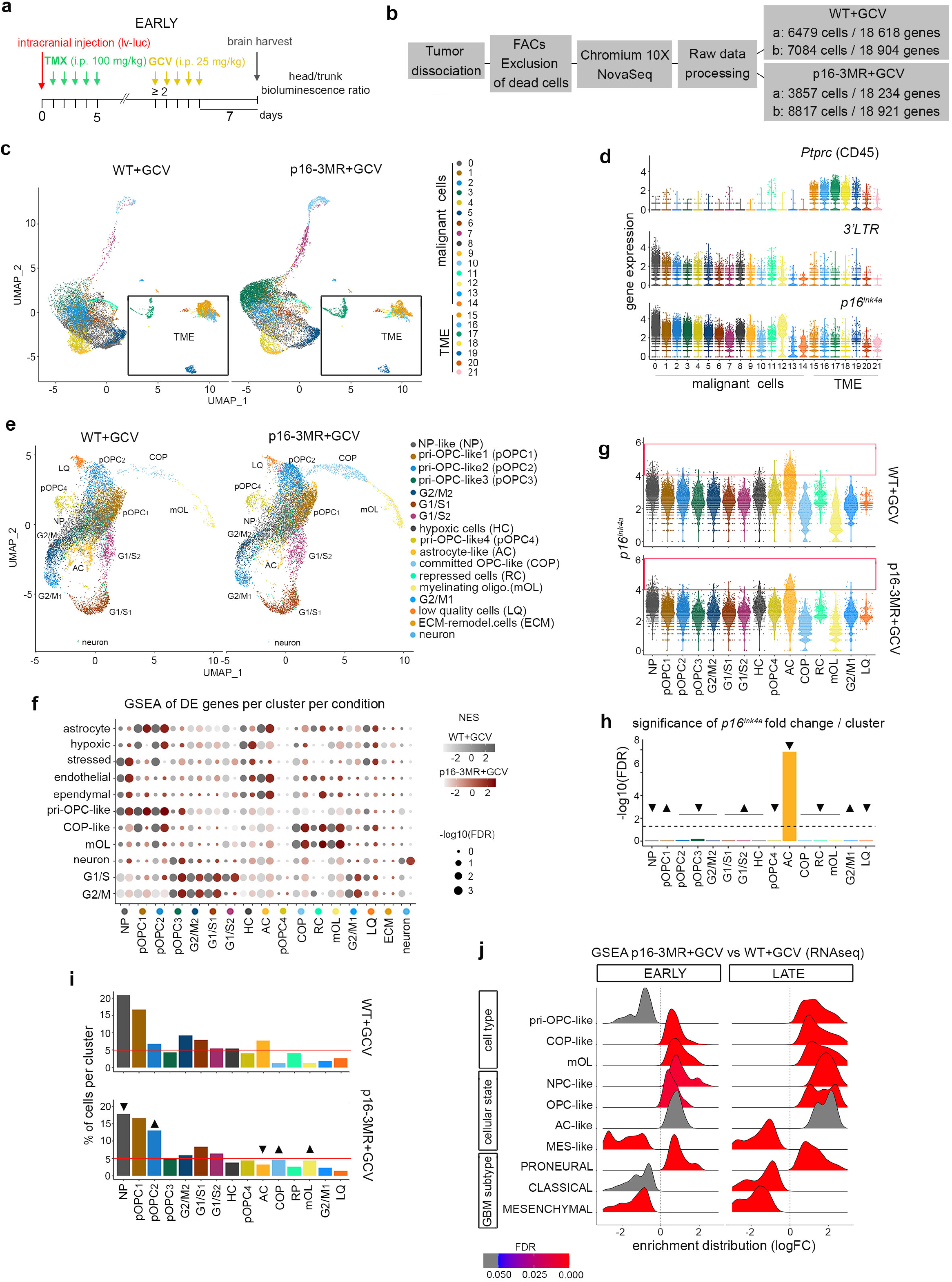
Identification of p16^Ink4a Hi^ cells in a subset of malignant cells. (**a**) Timeline of the mouse GBM generation for scRNAseq at the early timepoint. (**b**) Scheme of the scRNAseq experiment. (**c**) UMAP plots of WT+GCV (n=2) and p16-3MR+GCV (n=2) GBM cells at a 0.5 resolution and annotated malignant cells and TME cells. (**d**) Violin plots of the expression of *CD45*, 3’*LTR* and *p16^Ink4a^* in WT+GCV GBM cells per cluster. (**e**) UMAP plots of WT+GCV (n=2) and p16-3MR+GCV (n=2) GBM malignant cells and annotated cell type at a 0.6 resolution. (**f**) GSEA dot plots of DE genes (FDR<0.05; avlogFC>0.25) in WT+GCV (grey dots) and p16-3MR+GCV (red dots) GBMs of gene lists from Weng *et al*.^42^ (Supplementary Table 1). (**g**) Violin plots of the expression of *p16^Ink4a^* in malignant cells per cluster. The red box indicates the cells with an expression of *p16^Ink4a^* ≥ 4 (p16^Ink4a Hi^ cells). (**h**) Bar plots representing the significance of *p16^Ink4a^* fold change per cluster in p16-3MR+GCV GBMs compared with WT+GCV GBMs. The arrow heads point to a decrease (arrow heads down) of increase (arrow heads up) in the fold change. (**i**) Bar plots representing the percentage of malignant cells per cluster in WT+GCV and p16-3MR+GCV GBMs. The arrow heads point to clusters which cell number varies between the two conditions. (**j**) GSEA ridge plot of gene lists from Weng *et al*.^85^, Neftel *et al*.^23^, and Wang *et al*.^29^ (Supplementary Table 1) between p16-3MR+GCV and WT+GCV GBMs at the early and late time points. Analysis performed from bulk RNAseq data. TMX: tamoxifen; GCV: ganciclovir; vhc: vehicle; i.p.: intraperitoneal; lv-luc: lentivirus-luciferase; TME: tumor microenvironment; UMAP: uniform manifold approximation and projection; LTR: long terminal repeat; DE: differentially expressed; GSEA: gene set enrichment analysis; FDR: false discovery rate; NES: normalized enrichment score; r. enrichment score: running enrichment score.

These results prompted us to focus our analyses on the malignant cell compartment. The p16^Ink4a Hi^ senescent cells were mostly present in cluster 0 which comprises the highest cell number in WT+GCV GBMs (2910 out of 13 563 cells; Fig. 3d). Further UMAP clustering of malignant cells at 0.6 resolution identified 17 clusters in the two conditions (Fig. 3e). GSEA using the mouse gene lists published by Weng *et al*.^42^ allowed the malignant cell clusters to be assigned cellular identities, which predominantly included cycling cells, pri-oligodendrocyte progenitor cell-like (pri-OPC-like), committed OPC-like (COP-like), myelinating oligodendrocyte (mOL), astrocyte-like (AC), neural progenitor-like (NP-like), and hypoxic cells (HC) (Fig. 3e and f, Supplementary Fig. 3f; Supplementary Table 1). The labeling of the clusters was also in agreement with GSEA using human gene lists published by Bhaduri *et al*.^26^ (Supplementary Fig. 3g). Some clusters exhibited mixed cell identities. The astrocyte cluster shared gene signatures of astrocytes, endothelial cells and ependymal cells whereas the pri-OPC-like 1 (pOPC1) and pri-OPC-like 2 (pOPC2) clusters shared gene signatures of pri-OPC-like cells, astrocytes and COP cells (Fig. 3f). Of note, the enrichment score of each subpopulation differed very little between p16-3MR+GCV and WT+GCV GBMs, except for the pOPC1-3 clusters (Fig. 3f).

The p16^Ink4a Hi^ cells were mainly grouped in the astrocyte cluster and to a lesser extent in the NP-like cluster (Fig. 3g). The levels of *p16^Ink4a^* decreased significantly in the astrocyte cluster in p16-3MR+GCV GBMs compared with WT+GCV GBMs. No other clusters showed a significant difference in *p16^Ink4a^* levels between the two conditions (Fig. 3h). Therefore, this analysis identifies the astrocyte cluster as senescent. On line with a senescent phenotype, the astrocyte cluster shared an inflammatory signature (gene signatures of microglia and tumor associated macrophages) (Supplementary Fig. 3g). Remarkably, the percentage of cells in the astrocyte cluster decreased in p16-3MR+GCV GBMs compared with WT+GCV GBMs (astrocyte cluster from 7.75% to 3.21%; Fig. 3i), in agreement with the partial removal of p16^Ink4a Hi^ cells by the p16-3MR transgene in the presence of GCV.

Altogether, scRNAseq analysis identifies a cluster of senescent malignant cells displaying astrocytic and inflammatory phenotype.

### Partial removal of p16^Ink4a Hi^ malignant cells impacts the remaining malignant cells

We next analyzed whether the partial removal of p16^Ink4a Hi^ cells impacted the remaining malignant cells in our GBM model. Although the size of the tumors at this early timepoint differed between p16-3MR+GCV and WT+GCV GBMs (Supplementary Fig. a-c), the percentage of cycling cells (pOPC3, G2/M1, G2/M2, G1/S1, G2/S2 clusters) remained stable (WT+GCV: 28.78%; p16-3MR+GCV: 28.05%). In addition, three clusters of the oligodendroglial lineage pOPC2, COP and mOL, increased in cell proportions upon the partial removal of p16^Ink4a Hi^ cells (pOPC2 from 6.81% to 13.10%; COP from 1.24% to 4.60%; mOL from 1.31% to 4.30%; Fig. 3i). We further validated these results using bulk RNAseq data of WT+GCV and p16-3MR+GCV GBMs collected at the early timepoint (Supplementary Fig. 3h). GSEA using the Weng *et al*.^42^ gene lists showed no difference in the expression of cycling genes (Supplementary Fig. 3i). In contrast, there was an increase of transcripts associated with COP and mOL gene signatures upon the partial removal of p16^Ink4a Hi^ cells (Fig. 3i). The increase in cell numbers (scRNAseq) and in gene signatures (bulk RNAseq) of the oligodendroglial lineage suggests a shift of the malignant cellular states upon senolytic treatment. Indeed, GSEA revealed an increase in OPC-like and NPC-like states and their associated proneural transcriptional subtype following the partial removal of p16^Ink4a Hi^ cells. In parallel, GSEA showed a decrease in the MES-like state and the mesenchymal transcriptional subtype (Fig. 3j). Remarkably, all of these phenotypic traits perdured until the late timepoint (Fig. 3j).

Altogether, based on scRNAseq analyses, p16^Ink4a Hi^ senescent cells are a small subset of malignant cells. Their partial removal impacts the remaining malignant cells leading to a long-lasting switch to a more oligodendroglial-like phenotype and a decrease in expression of genes signatory of a mesenchymal cell identity.

### Modulation of the immune compartment following the partial removal of p16^Ink4a Hi^ cells

The mesenchymal transcriptional GBM subtype is associated with enhanced expression of anti-inflammatory and tumor-promoting macrophages^29,31,43^. We therefore examined the immune compartment in the scRNAseq data at the early timepoint following the partial removal of p16^Ink4a Hi^ cells. UMAP clustering of *CD45*+ cells revealed seven clusters in the two experimental conditions (Fig. 3d, Supplementary Fig. 3d and Fig. 4a). Differentially expressed (DE) genes and GSEA allowed the labelling of these clusters into infiltrating bone marrow-derived macrophages (BMDM), resident microglia and T cells^44^ (Fig. 4b and c, Supplementary Fig. 4a; Supplementary Table 1). All the BMDM and microglia clusters harbored an antiinflammatory gene signature. Furthermore, the BMDM-like1 and microglia clusters shared an antagonist pro-inflammatory gene signature^32^ (Fig. 4c; Supplementary Table 1). In addition, the proportion of the immune fraction within the tumor hardly varied between WT+GCV and p16-3MR+GCV GBMs. However, the number of T cells increased (from 9% to 27%), whereas the number of BMDM decreased (from 41% to 30%) upon the partial removal of p16^Ink4a Hi^ cells (Fig. 4d). This latter phenotype was confirmed by GSEA on bulk RNAseq data, which showed a significant decrease in transcripts associated with a core BMDM signature at the early and late timepoints in p16-3MR+GCV GBMs compared with controls (Supplementary Fig. 4b). In addition, an estimation of the abundances of the main immune cell types from our bulk RNAseq data using CIBERSORT pointed to a significant decrease of BMDM upon partial removal of p16^Ink4a Hi^ cells at the late timepoint (Supplementary Fig. 4c).

**Figure 4.**
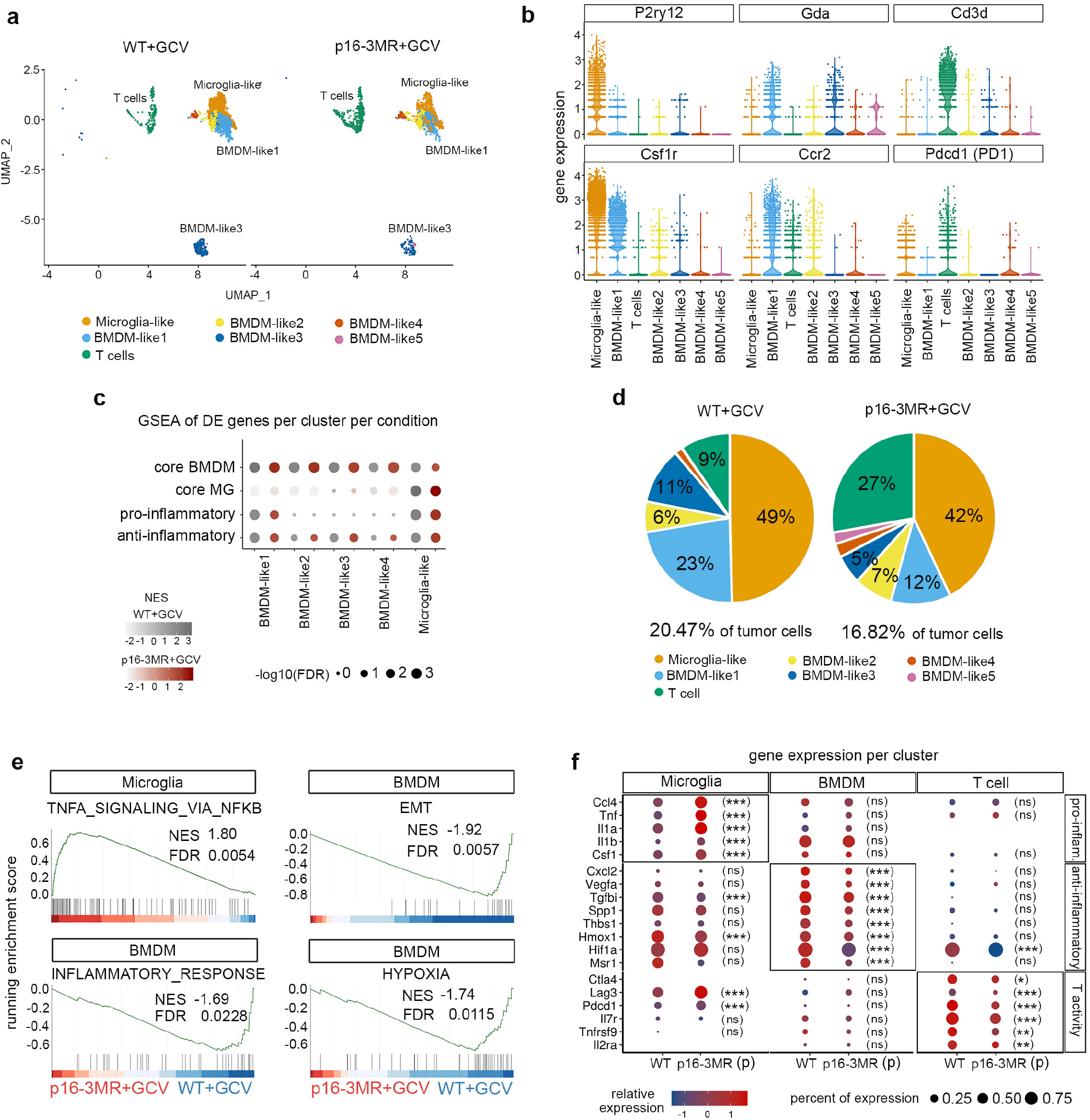
Modulation of the immune compartment following p16^Ink4a Hi^ cells partial removal. (**a**) UMAP plots of *CD45+* cells in WT+GCV and p16-3MR+GCV GBMs at a 0.5 resolution and annotated cell type. (**b**) Violin plots representing the expression of selected DE genes (FDR<0.05; avlogFC>0.25) per cluster in WT+GCV GBMs. (**c**) GSEA dot plot of DE genes (FDR<0.05; avlogFC>0.25) in WT+GCV (grey dots) and p16-3MR+GCV (red dots) *CD45+* clusters of core-BMDM, core-MG, pro-inflammatory and antiinflammatory pathways as defined in Bowman *et al*.^44^ and Darmanis *et al*.^32^ (Supplementary Table 1). (**d**) Chart pies representing the percentage of *CD45*+ cells per cluster in WT+GCV and p16-3MR+GCV GBMs. (**e**) GSEA graphs representing the enrichment score of Hallmark gene lists in p16-3MR+GCV compared with WT+GCV microglia clusters and pooled BMDM clusters. The barcode plot indicates the position of the genes in each gene set; red represents positive Pearson’s correlation with p16-3MR+GCV expression, blue with WT+GCV expression. (**f**) Dot plots of the relative expression of selected genes in WT+GCV and p16-3MR+GCV microglia, pooled BMDM and T cells clusters. Statistical significance of the expression of genes in p16-3MR+GCV compared with WT+GCV clusters was determined by Wilcoxon-Mann-Whitney test (ns, not significant, *, p<0.05; **, p<0.01; ***, p<0.001). UMAP: uniform manifold approximation and projection, BMDM: bone marrow-derived macrophages; MG: microglia; DE: differentially expressed; EMT: epithelial to mesenchymal transition; GSEA: gene set enrichment analysis; FDR: false discovery rate; NES: normalized enrichment score.

We next examined whether the activity of immune cell types was altered in GBM tumors partially depleted of senescent cells. GSEA on scRNAseq data at the early timepoint revealed an upregulation of TNFA signaling via the NFKB pathway in the microglia cluster and a downregulation of genes associated with the epithelial-to-mesenchymal transition, inflammatory and hypoxia pathways in the BMDM clusters in p16-3MR GBMs compared with WT+GCV GBMs (Fig. 4e). Close examination of the DE genes in these pathways revealed a significant increase in the expression of genes associated with a pro-inflammatory signature (*Ccl4, Tnf, Il1a, Il1b, Csf1*) in the microglia cluster and a significant decrease in the expression of genes related to an anti-inflammatory signature (*Cxcl2, Vegfa, Tgfbi, Spp1, Thbs1, Hmox1, Hif1a*) in the BMDM clusters (Fig. 4f, Supplementary Table 2). In addition, T cell cluster analysis revealed a decrease in the expression of genes regulating the activity of T cells, including the immune checkpoint genes, *Ctla4, Lag3* and *Pdcd1* (encoding PD1) (Fig. 4f). Consistent with these data, GSEA of bulk RNAseq data revealed a decrease in transcripts associated with an anti-inflammatory pathway following the partial removal of p16^Ink4a Hi^ cells at the early and late timepoints (Supplementary Fig. 4b).

Collectively, the transcriptomic analysis at single cell and bulk levels shows that the partial removal of p16^Ink4a Hi^ malignant cells modulates the abundance and the activity of tumor associated macrophages.

### Identification of NRF2 activity and its putative targets in p16^Ink4a Hi^ malignant cells

To explore the regulators of senescence in p16^Ink4aHi^ malignant cells, we performed pathway enrichment analysis with the ENCODE and ChEA consensus TFs from ChIP-X database using Enrichr^45^ on three gene sets enriched in p16^Ink4a Hi^ senescent cells: (i) differentially downregulated genes in the p16-3MR+GCV vs the WT+GCV astrocyte cluster from the scRNAseq data at the early timepoint (Early; Fig. 5a and b; Supplementary Table 3); (ii) differentially upregulated genes in p16^Ink4a^ positive vs p16^Ink4a^ negative malignant cells from scRNAseq analysis at the late timepoint (Late (1); Fig. 5c and d, Supplementary Fig. 5a-e; Supplementary Table 3); (iii) differentially downregulated genes in the p16-3MR+GCV vs the WT+GCV GBMs from the bulk RNAseq data at the late timepoint (Late (2); Fig. 5e and f; Supplementary Table 3). Remarkably, the *Nuclear Factor Erythroid 2 Like 2* (*Nfe2l2*) signaling pathway was enriched in the three gene sets. NRF2 encoded by *Nfe2l2* is an antioxidant defense system that appears to be a plausible candidate to trigger the pro-tumoral action of p16^Ink4a Hi^ senescent cells as it induces cellular senescence in fibroblasts^46^ and confers a selective advantage in cancer cells^47^. Among the identified NRF2 putative targets, three genes were common to all three data sets (*Tgif1, Plaur, Gja1*) whereas eight genes were shared between two of the three gene lists (*Dap, Esd, Lmna, Areg, Igfbp3, Cdkn2b, Tnc* and *Peak1*) (Fig. 5g; Supplementary Table 3). As illustrated on the heatmap, the combined expression of *Nfe2l2* and 11 putative target genes were unique to p16^Ink4a Hi^ cells in WT+GCV GBMs (Fig. 5h).

**Figure 5.**
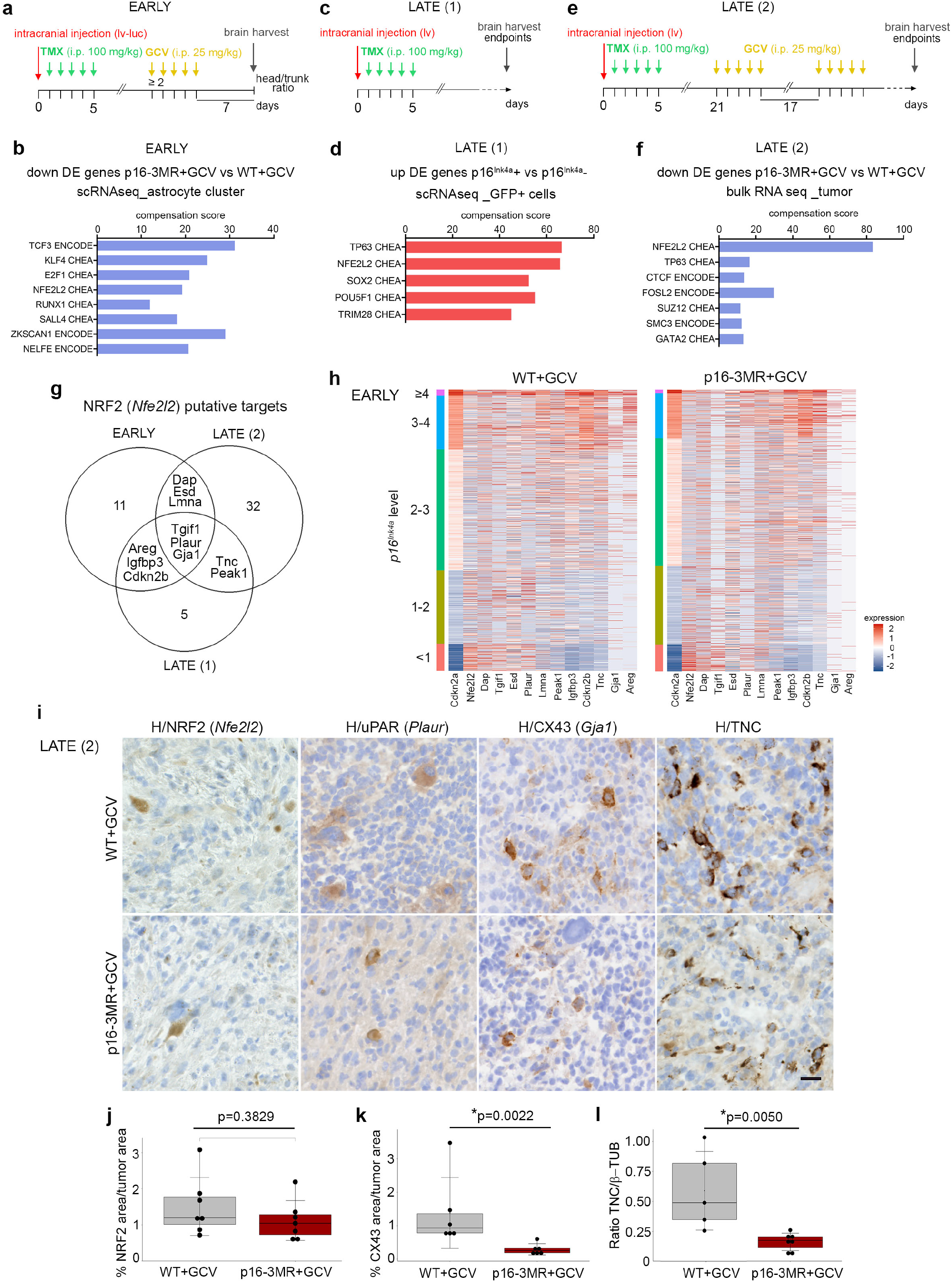
Identification of NRF2 activity and its putative targets in p16^Ink4a Hi^ malignant cells. (**a**) Timeline of the mouse GBM generation for scRNAseq at the early timepoint (EARLY). (**b**) Barplot corresponding to significantly enriched pathways (ENCODE and ChEA consensus TFs from ChIP-X, Enrichr) in differentially downregulated genes (FDR<0.05; avlogFC>0.25) in the p16-3MR+GCV compared with the WT+GCV astrocyte clusters from the scRNAseq data (as shown in **a**). (**c**) Timeline of the mouse GBM generation for scRNAseq at the late timepoint (LATE (1)). (**d**) Barplot corresponding to significantly enriched pathways in differentially up-regulated genes (FDR<0.05; logFC>0.5) in p16^Ink4a^ positive vs p16^Ink4a^ negative malignant cells from the scRNAseq data (as shown in **c**). (**e**) Timeline of the mouse GBM generation for bulk RNAseq at the late timepoint (LATE (2)). (**f**) Barplot corresponding to significantly enriched pathways in differentially down-regulated genes (FDR<0.05; logFC>0.5) in p16-3MR+GCV compared with WT+GCV GBMs from the bulk RNAseq data (as shown in **e**). (**g**) Venn diagram of NRF2 putative targets between the 3 gene sets as shown in **a**, **c** and **e**. (**h**) Heatmaps of *Nrf2* and its 11 identified putative targets in WT+GCV and p16-3MR GBMs. Cells are classified in 5 categories according to *p16^Ink4a^* expression levels. (**i**) Representative immunohistochemistry (IHC, brown) counterstained with hematoxylin (H, purple) on mouse GBM cryosections at the late timepoint. H: hematoxylin. (NRF2: WT+GCV, n=7; p16-3MR+GCV n=7; uPAR: WT+GCV, n=3; p16-3MR+GCV, n=3; CX43: WT+GCV, n=5; p16-3MR+GCV, n=6; TNC: WT+GCV, n=5; p16-3MR+GCV, n=7 independent mouse GBMs). Scale bar: 20 μm. (**j**) Quantification of the NRF2 area (IHC) over the tumor area (WT+GCV, n=7; p16-3MR+GCV, n=7 independent mouse GBMs). (**k**) Quantification of the CX43 area (IHC) over the tumor area (WT+GCV, n=5; p16-3MR+GCV, n=6 independent mouse GBMs). (**l**) Quantification of the ratio of TNC over β-TUBULIN expression (western blot) (WT+GCV, n=5; p16-3MR+GCV, n=7 independent mouse GBMs). Raw data are shown in Supplementary Fig. 5g. **j**, **k**, **l**: data are presented as the mean ± SD. Statistical significance was determined by Wilcoxon-Mann-Whitney test (*, p<0.05). i.p.: intraperitoneal; lv: lentivirus; lv-luc: lentivirus-luciferase; TMX: tamoxifen; DE: differentially expressed; GCV: ganciclovir; H: hematoxylin;

Immunohistochemistry on GBM cryosections collected at the late timepoint revealed that NRF2 was expressed in a few scattered cells (Fig. 5i and Supplementary Fig. 5f). Quantification of the NRF2 expression area in the tumor showed a modest decrease in p16-3MR+GCV compared with WT+GCV GBMs (Fig. 5j). Of note, the expression of NRF2 in cells expressing low levels of *p16^Ink4a^*, most probably CD45+ cells, may have concealed decreased NRF2 expression in senescent cells (Fig. 5h). We further examined the expression of three NRF2 putative target genes whose encoded proteins are associated with senescence, glioma progression or glioma resistance, respectively, namely urokinase plasminogen activator receptor (uPAR) encoded by *Plaur*^48^, Tenascin-C (TNC)^49^ or Connexin43 (CX43) encoded by *Gja1*)^50^. These proteins were expressed in a few scattered cells throughout the tumor. TNC was expressed in more cells than uPAR and CX43 in line with their transcript expression at the single cell level (Fig. 5h and i, Supplementary Fig. 5f). Quantification of CX43 by IHC and TNC by western blot revealed a significant downregulation of these proteins in p16-3MR+GCV GBMs compared with WT+GCV GBMs, strengthening *Gja1* and *Tnc* as NRF2 target genes in GBM (Fig. 5k and l, Supplementary Fig. 5g). We then assessed whether interactions between NRF2 selected targets and the immune fractions were observed in GBMs. We interrogated for ligand-receptor interactions between cluster 0, enriched in p16^Ink4a Hi^ cells, and the immune clusters in the scRNAseq data at the early timepoint using CellPhoneDB (Fig. 3d, Supplementary Fig. 5h). *In silico* analysis highlighted possible interactions between TNC and PLAUR, expressed in malignant cells, and integrins receptors expressed in the immune clusters. Remarkably, putative TNC-aVb3 and PLAUR-aVb3 ligand-receptor interactions between malignant cells and T cells were abolished upon partial removal of p16^Ink4a Hi^ cells (Supplementary Fig. 5h).

All together these data identify NRF2 activity and its putative target genes in p16^Ink4a Hi^ senescent cells and suggest that this signal could in part trigger the detrimental action of senescent cells during gliomagenesis.

### Knockdown of NRF2 in malignant cells recapitulates most features of the senolytic treatment

NRF2 has pleiotropic actions depending on cellular context. Tumor suppressing effects of NRF2 are mediated via the maintenance of a functional immune system^47^. For instance, in a mouse lung cancer model, NRF2 activity in immune cells contributes to suppress tumor progression^51^. To directly test whether NRF2 triggers the tumor promoting action of malignant senescent cells, we used a knockdown approach, introducing a microRNA targeting NRF2 (miR-NRF2) into the lentivirus used to induce gliomagenesis. We analyzed the resultant tumors at the late timepoint (Fig. 6a and b, Supplementary, Fig. 6a). Quantification of NRF2 by IHC revealed a significant decreased of the protein in miR-NRF2-GBMs compared with miR-control (ctl)-GBMs (Fig. 6c and d, Supplementary Fig. 6d). Notably, NRF2 is also expressed in CD45+ cells, not targeted by our approach, which persisted in miR-NRF2-GBMs. We performed bulk RNAseq and GSEA of miR-NRF2- and miR-ctl-GBMs at late timepoint, and found a significant downregulation of canonical NRF2 targets and NRF2 targets from the combined analysis, confirming knockdown of NRF2 using our miR-NRF2 (Fig. 6e; Supplementary Table 3).

**Figure 6.**
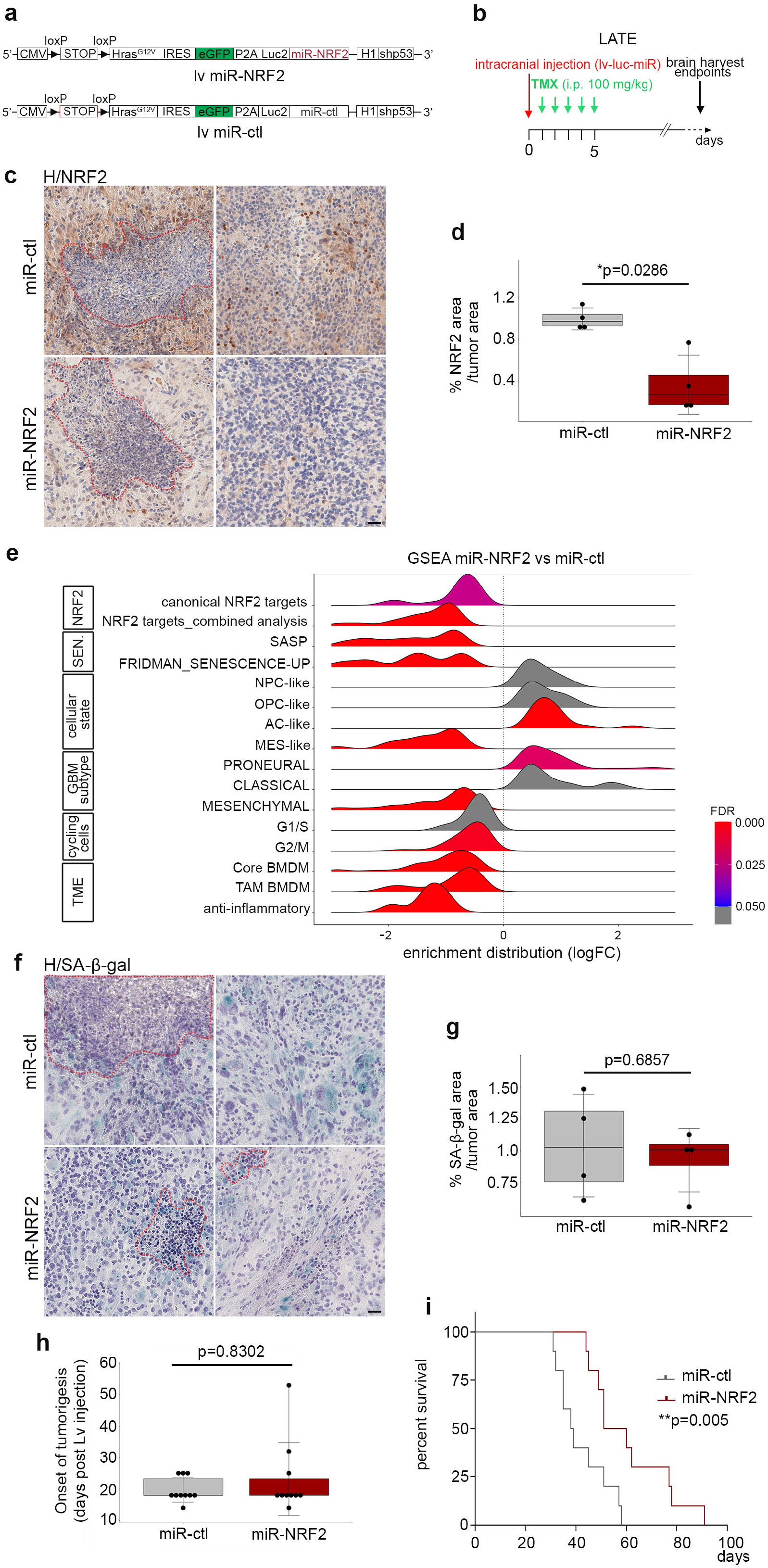
Knockdown of NRF2 in malignant cells recapitulates most features of the senolytic treatment. (**a**) Scheme of the lentiviral vector containing either a miR-NRF2 or a miR-ctl. (**b**) Timeline of the mouse GBM generation at the late timepoint. (**c**) Representative NRF2 IHC staining (brown) on miR-ctl (n=4) and miR-NRF2 (n=4) GBM cryosections. Necrotic areas are outlined in red dashed lines. (**d**) Quantification of the NRF2 area (IHC) over the tumor area (miR-ctl n=4; miR-NRF2 n=4). (**e**) GSEA ridge plot on bulk RNAseq of miR-NRF2-GBMs compared with miR-ctl-GBMs (see Supplementary Table 1 for gene lists). (**f**) Representative SA-β-gal (blue) staining on miR-ctl (n=4) and miR-NRF2 (n=4) GBM cryosections. Necrotic areas are outlined in red dashed lines. (**g**) Quantification of the SA-β-gal area over the tumor area (miR-ctl n=4; miR-NRF2 n=4). (**h**) Boxplot representing the onset of tumorigenesis in miR-ctl (n=10) and miR-NRF2 (n=10) mice following post-lentiviral injection. The onset of tumorigenesis is defined when the bioluminescence reached 3e10^6^. (**i**) Kaplan-Meier survival curves of miR-ctl (n=10, median survival 38.5 days) and miR-NRF2 mice (n=10, median survival 55.5 days). Statistical significance was determined by Mantel-Cox log-rank test (**, p<0.01). Scale bar, **c** and **f**: 50 μm. **d**, **g, h**: Statistical significance was determined by Wilcoxon-Mann-Whitney test (*, p<0.05). lv: lentivirus; miR-ctl: miR-control; H: hematoxylin; GSEA: gene set enrichment analysis; sen.: senescence; TAM: associated macrophages; BMDM: bone marrow derived macrophages.

We next asked whether knockdown of NRF2 in malignant cells impacted cellular senescence. The percentage of the tumor area encompassing SA-β-gal cells was similar in miR-NRF2 GBMs compared with miR-ctl GBMs, suggesting that NRF2 knockdown in malignant cells does not induce the death of senescent cells (Fig. 6f and g, Supplementary Fig. 6e). However, GSEA performed on bulk RNAseq data from miRNRF2- and miR-ctl-GBMs revealed a significant downregulation of SASP genes associated with senescence (Fig. 6e). Among these genes, *Mmp1a, Mmp3, Mmp10, Plau, Col1a2, Timp1* and *Thbs1* were differentially expressed between the two conditions (Supplementary Fig. 6b). This result strongly suggests that NRF2 regulates directly or indirectly the expression of SASP genes. Further, we examined whether NRF2 knockdown mimicked the phenotype of senolytic treatment. GSEA revealed a significant decrease in the expression of genes associated with a mesenchymal identity, a BMDM signature and anti-inflammatory pathways, similar to the gene signature changes observed in p16-3MR+GCV GBMs (Fig. 6e; Supplementary Fig. 6c). However, genes associated with an oligodendroglial identity were not modulated upon NRF2 knockdown (Fig. 6e), in contrast to the effect of the partial removal of p16^Ink4a Hi^ senescent cells.

Finally, as NRF2 activity protects against DNA-damaging agents and prevents carcinogenesis^52^, we explored whether NRF2 knockdown impacted the onset of tumorigenesis. Live bioluminescence imaging showed no difference in the onset of tumorigenesis between the two tumor types (Fig. 6h). Most importantly, the presence of miR-NRF2 in malignant cells significantly increased the survival of GBM-bearing mice compared with controls (Fig. 6i), an effect that was more marked than upon the partial removal of p16^Ink4a Hi^ senescent cells (Fig. 2c and d; Fig. 6i). One major difference between the two paradigms was the decrease of transcripts linked to cell cycle, which occurred only in the knockdown of NRF2 in malignant cells (Fig. 6e and Supplementary 6c). One reason for this difference could be that all malignant cells were targeted by the miR strategy, whereas the p16-3MR+GCV paradigm only partially removed senescent cells.

Collectively our results show that NRF2 knockdown recapitulates most features of senolytic treatment and strongly support NRF2 as a cellular senescence regulator in malignant cells.

### Mouse senescent signature is conserved in patient GBMs and its enrichment score is predictive of a worse survival

We next explored whether p16^Ink4a Hi^ senescent cells are conserved in patient GBMs. We first established a senescent signature from scRNAseq data at the early timepoint (Fig. 3). We compared the transcriptome of p16^Ink4a Hi^ cells in astrocyte and NP-like clusters (p16^Ink4a Hi^ group) with the remaining malignant cells in WT+GCV GBMs (Fig. 3g, Fig. 7a and b). GSEA in the p16^Ink4a Hi^ group revealed a downregulation of cell cycle pathways and an oligodendroglial state, and increased expression of genes associated with inflammation, NRF2 signaling, MES-like state, mesenchymal transcriptional GBM subtype (Supplementary Fig. 7a and b). We further selected a list of 31 genes to define a GBM senescence signature. Among the 278 differentially upregulated genes (FDR<0.05) in the p16^Ink4a Hi^ group, we selected genes that were expressed in more than 90% of p16^Ink4a Hi^ cells and presented a log2-fold change superior to 0.8 between the two groups. As expected, senescence-associated genes were enriched in the astrocyte cluster and to a lesser extent in the NP-like cluster (Fig. 7c). The encoded proteins were associated with diverse cellular processes compatible with cellular senescence, such as cell cycle arrest (*Cdkn1a, Cdkn2a, Cdkn2b*), lysosomal function (*Ctsb*, *Ctsd, Ctsl, Ctsz, Lamp1, Lamp2*), cellular growth (*Igfbp2, Igfbp3*), extracellular matrix interaction (*Sparc, Tnc, Sdc4, Lgals1, Timp1, Mt1*), cytoskeleton interaction (*Pdlim4, S100a11, Tmsb4x, Sep11*) and cancer (*Tm4sf1, Ociad2, Emp3*) (Fig. 7c). We then computed a single-sample GSEA (ssGSEA) senescent Z-score corresponding to the enrichment Z-score of the 31 genes of the senescence signature in all malignant cells of WT+GCV and p16-3MR+GCV GBM transcriptomes. As expected, the astrocyte cluster in WT+GCV GBMs contained the largest high senescent Z-score distribution rate (Supplementary Fig. 7c). For unbiased analysis, we defined the high distribution rate as the highest decile. This percentage dropped in all clusters in p16-3MR+GCV GBMs compared with WT+GCV GBMs, as predicted by the partial removal of p16^Ink4a Hi^ senescent cells (Supplementary Fig. 7c).

**Figure 7.**
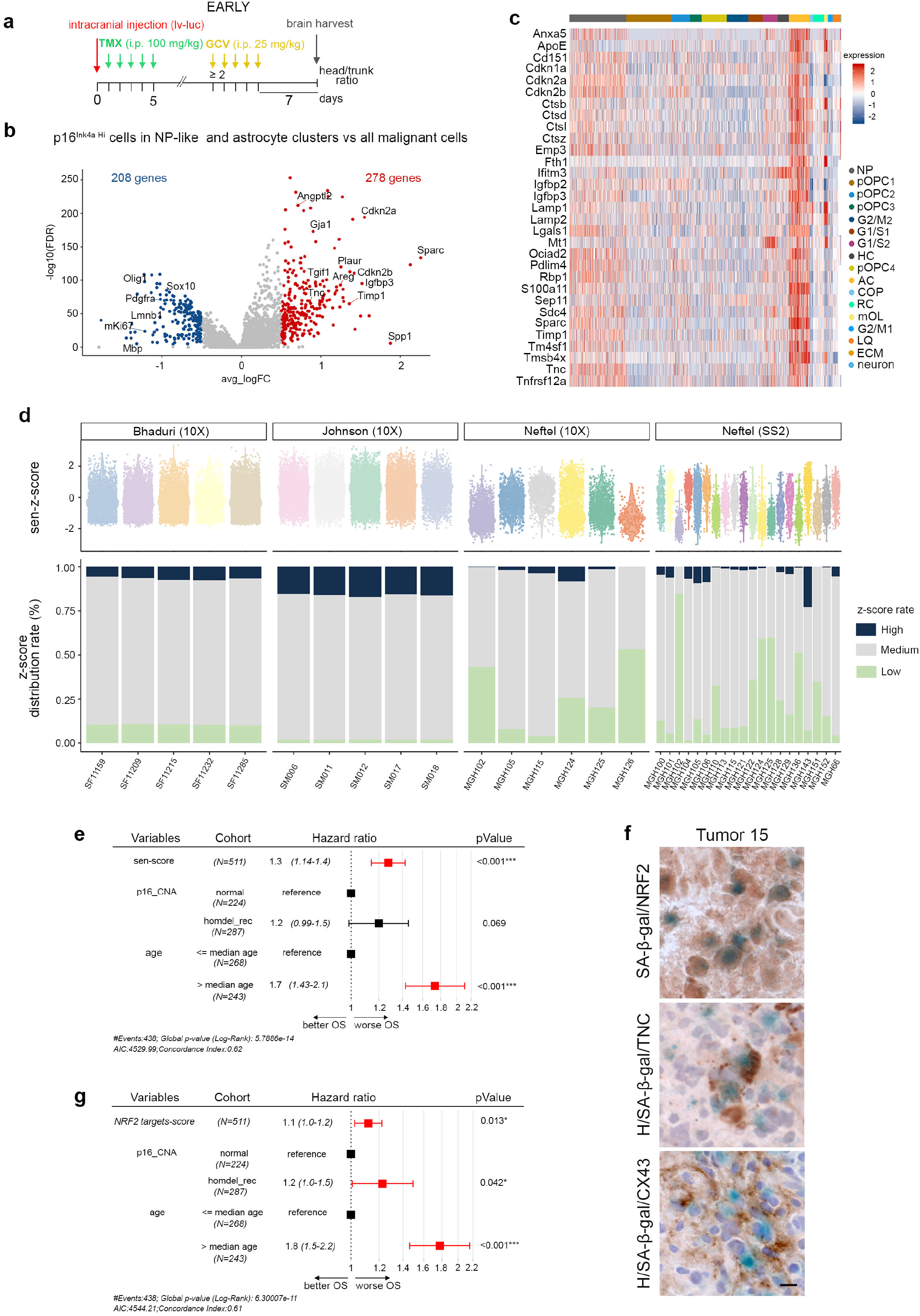
Mouse senescent signature is conserved in patient GBM and its enrichment score is predictive of a worse survival. (**a**) Timeline of the mouse GBM generation for scRNAseq at the early timepoint. (**b**) Volcano plot of differentially expressed (DE) genes (−0.5<log2FC>0.5; FDR<0.05) between p16^Ink4a Hi^ cells (gene expression ≥ 4) of astrocyte and NP-like clusters compared with the remaining malignant cells in WT+GCV GBMs. (**c**) Heatmap of the 31 senescence signature genes in WT+GCV GBMs. (**d**) Top: Violin plots of the single-sample GSEA (ssGSEA) senescent Z-score in all patient GBM cells. Patient GBMs data were extracted from Bhadury *et al*.^26^, Johnson *et al*.^53^ and Neftel *et al*.^23^. Bottom: Barplots of the percentage of the ssGSEA senescent Z-score distribution rate in all patient GBM cells. High and Low distribution rates correspond to the highest and lowest decile, respectively. (**e**) Table representing a Cox regression model using the ssGSEA-senescence score (sen-score), p16^INK4a^ copy number alteration (p16-CNA) stratified into a group without alteration (normal) and a group harboring homozygous recessive deletion (homdel_rec) in the *INK4a* locus and the age of the patients. (**f**) Representative SA-β-gal staining (blue) coupled with IHC (brown) and counterstained with hematoxylin (H) on patient GBM cryosections. 3 patient GBMs were analyzed per antibody. Scale bar :10 μm. (**g**) Table representing a Cox regression model using the ssGSEA NRF2 targets score (NRF2 targets score), p16^INK4a^ copy number alteration and the age of the patients. **e** and **g**: data were extracted from The Cancer Genome Atlas (TCGA) GBM data sets. GCV: ganciclovir; TMX: tamoxifen; i.p.: intraperitoneal; lv: lentivirus; lv-luc: lentivirus-luciferase; OS: overall survival.

We then applied the ssGSEA senescence Z-score to three single cell data sets of patient GBMs^23,26,53^. We analyzed separately the Neftel dataset according to the sequencing technology (Smartseq2 (SS2) and 10X). The range of the ssGSEA senescence Z-score of cells from patient GBMs was similar to those identified in mouse GBMs (Fig. 7d; Supplementary Fig. 7c). Hence, similar senescent transcriptomic profiles were observed in cells from mouse and patient GBMs. GBMs from the three data sets contained high senescent Z-score distribution rates, with the exception of 2/31 tumors, possibly due to the small number of cells sequenced in these samples (MGH126: 201 cells and MGH151: 151 cells). In summary, this analysis strongly suggest that the mouse senescent signature is conserved in cells from patient GBMs.

To assess whether the ssGSEA senescence score could be used as a prognostic factor for patients with GBM, we interrogated The Cancer Genome Atlas (TCGA) GBM data sets and performed a Cox regression analysis with three variables. Cellular senescence was linked to aging and our mouse senescence signature was based on the expression of p16^Inka^. Therefore, we used as variables the ssGSEA senescence score, p16^Ink4a^ copy number alteration status and the age of the patient. The Cox regression model showed that regardless of p16^Ink4a^ status and the age of the patient, the enrichment of the senescence score predicted a worse survival (hazard ratio above 1) in patients with GBM (Fig. 7e).

Finally, we tested whether NRF2 activity could account for some of the tumor promoting action of cellular senescence in patient GBMs, similar to mouse GBMs. First, SA-β-gal staining coupled with IHC on cryosections revealed the expression of NRF2, TNC and CX43 in SA-β-gal+ cells in patient GBM samples (Fig. 7f and Supplementary 7d). As described above, we next interrogated ssGSEA NRF2 target scores in TCGA GBM data sets and performed a Cox regression analysis. NRF2 targets corresponded to the 59 genes identified in the combined analysis (Supplementary Table 3; Fig. 5g). The Cox regression model showed that regardless of p16^Ink4a^ status and the age of the patient, an enriched NRF2 target gene score predicted worse survival in patients with GBM (Fig. 7g).

In summary, our data show that cells enriched for the mouse senescent signature are present in patient GBMs and that the enrichment scores of senescence and of NRF2 targets are correlated with a worse survival in patients with GBM.

## Discussion

Depending on the context, cellular senescence plays both beneficial and detrimental roles during tumor progression. Here, we revealed the tumor promoting action of malignant senescent cells in mouse and patient GBMs. The mouse MES-GBM model used in the present study, even though its genetic differs from the patient GBMs, recapitulated the histopathology, the heterogeneity of cellular states, the infiltration of BMDM specific to the mesenchymal transcriptional GBM subtype and the senescent features of patient GBMs (see also^43^). Partial removal of p16^Inka Hi^ malignant senescent cells modified the tumor ecosystem and improved the survival of GBM-bearing mice. The difference of survival following a senolytic treatment appeared to be relatively modest, nonetheless this difference was significant and was observed in two replicates using the p16-3MR paradigm and in the ABT263 paradigm. This result is remarkable given the fact that senescent cells represented less than 7% of the tumor and that their removal using the p16-3MR transgene was only partial. These findings suggest that senolytic drug therapy may be a beneficial adjuvant therapy for patients with GBM.

By combining single cell and bulk RNA sequencing, immunohistochemistry and genetic knockdowns, our study established a link between senescence and NRF2 activity in the context of GBM. Previous work demonstrated that chronic activation of NRF2 contributes to tumor growth, metastasis, treatment resistance and poorer prognosis in patients with cancer^47^. NRF2 binds to antioxidant responsive elements (AREs) and controls the expression of a battery of genes regulating metabolism, intracellular redox-balance, apoptosis, and autophagy^47^. Depending on the context, NRF2 promotes or delays fibroblasts senescence^46,47^. *Nrf2* transcription can be induced by oncogenes (e.g. KRAS) and its activity is modulated by environmental cues (e.g. hypoxia, ROS)^54^. Under homeostatic state, cytoplasmic NRF2 binds to KEAP1, which mediates its proteasomal degradation. However, impairment of NRF2-KEAP1 binding, either by phosphorylated p62 or by elevated ROS permits NRF2 nuclear translocation and activation of target genes^55,56,57^. Previous studies on GBMs showed that the KEAP1-mediated degradation of NRF2 promotes *in vitro* glioma stem cell survival and that NRF2 is hyperactivated in the mesenchymal transcriptional GBM subtype^58^. In the present study, we identified NRF2 putative targets that are not canonical NRF2 targets. These targets encode for growth factors (AREG, IGFBP3, TGIF1), ECM components or remodelers (TNC, uPAR, ESD) or cell-cell interactors (CX43) and have been previously identified as SASP factors^39,40,41^. TNC and CX43 are of particular interest. Indeed, CX43 participates in the formation of microtubes that interconnect malignant cells, creating a cellular network resistant to treatment^50^. Further, the pro-tumoral functions of TNC has been described, independently of the senescence context^59,60^. TNC is a component of the glioma ECM that binds to integrin receptors, EGF receptor (EGFR), SYNDECAN 4 (SDC4) and regulates angiogenesis, proliferation and cell migration^49^. Hence, TNC functions could partly be responsible for the tumor promoting phenotype of the p16^Ink4a Hi^ malignant senescent cells. As mentioned above, NRF2 has pleiotropic actions depending on cellular context and is expressed in multiple cell types. Of note, in this study, we did not address the function of NRF2 in the GBM microenvironment, notably in CD45+ cells. To sum up, our findings suggest that a senolytic treatment may represent a novel therapeutical strategy to eliminate NRF2-malignant senescent cells without targeting cells from the microenvironment.

Single cell RNAseq analysis of mouse GBMs allowed the comprehensive characterization of the pro-tumorigenic malignant senescent cells. Although our approach focused primarily on p16^Ink4a Hi^ senescent cells in a mouse MES-GBM model, our findings show that the senescence signature we established in this study, is applicable to GBMs regardless of p16^Ink4a^ status (Fig. 7e). The presence in the senescence signature of three genes (*Cdkn2a, Cdkn2b, Cdkn1a*) encoding for cyclin dependent kinase inhibitors warrants cell cycle exit and entry into senescence. Of note, *CDKN1A* (*p21^CIP1^*) is rarely mutated in patient GBMs (0.4%) and p21^CIP1^ mediates senescence in many tissues^3^. Furthermore, we presume that the senescent signature defined in the study is specific to detrimental senescence. Indeed, the enrichment of the senescence score predicts a worse survival in patients with GBM and multiple genes in the signature encode for proteins which activities are associated with tumor aggressiveness and/or worse patient prognosis (CD151^61^; EMP3^62^; IGFBP2^63^; LGALS1^64^; TMSB4X^65^;TNC/SDC4^60^; SPARC^66^; TIMP1^67^). Future studies will determine whether the senescence scoring could be used in the diagnosis of patients with GBM to improve the design of personalized treatment and effective combinatorial strategies.

In this study we showed that senolytic treatments applied to GBM bearing mice delays temporarily tumor growth (Fig. 2c and d). These findings raise the question of the benefit of combining senotherapy to improve response to other therapies. The field of senotherapies, which includes drugs eliminating senescent cells (senolytic drugs: anti-BCL2 and BCL-xL, Dasatinib and Quercitin, cardiac glycosides) or drugs inhibiting their function (senostatic drugs such as Metformin) is under active investigation^68,69,70,71^. A recent study reported in a MC38 mouse syngeneic tumor model, that Venetoclax (anti-BCL2) augments the antitumor efficacy of an anti-PD1 treatment by increasing the T effector memory cells although Venetoclax reduces the overall T cell number, a known function of the anti-apoptotic protein inhibitors^15^. The authors reported the absence of cancer cell-intrinsic effects of the Venetoclax. In sharp contrast, our data provide evidence that the senolytic p16-3MR transgene in a mouse GBM model removes malignant senescent cells and decreases the anti-inflammatory phenotype. Although a thorough study on the consequence of senolytic treatment on the immune system is required, we would like to propose that senolytic treatment may prime GBM to respond to immunotherapy. This hypothesis is attractive as the immunotherapies with the anti-PD1 and PD-L1 antibodies did not show an extension of the overall survival in treating patients with recurrent GBM^72,73^. One possible explanation for this failure could be that GBMs contain very few immune effector cells^74^. Further work on immunocompetent GBM models is now needed to evaluate the effect of novel senolytic/senostatic treatments on gliomagenesis and assess their efficacy as companion therapy.

## Methods

### Patient samples

Fresh patient GBM samples were selected from the Pitié-Salpêtrière tumor bank Onconeurotek. They were reviewed by our senior pathologist (FB) to validate the histological features and confirm patients’ diagnosis. Collection of tumor samples and clinical-pathological information were obtained upon patients’ informed consent and ethical board approval, as stated by the Declaration of Helsinki. Molecular characterizations were performed as previously described^75^.

### Mouse and breeding

All animal care and treatment protocols complied with European legislation (no. 2010/63/UE) and national (French Ministry of Agriculture) guidelines for the use and ethical treatment of laboratory animals. All experiments on animals were approved by the local ethics committee (approval APAFIS 9131). To generate the GBM mouse model, we crossed Glast^creERT2/+^ mice^76^ with the Pten^fl/fl^ mice^77^. Glast^creERT2/+^; Pten^fl/fl^ males were bred with either Pten^fl/fl^ or Pten^fl/fl^; p16-3MR/+ females^17^ to generate Glast^creERT2/+^; Pten^fl/fl^ and Glast^creERT2/+^; Pten^fl/fl^; p16-3MR/+ mice, named WT and p16-3MR mice respectively. All animals used in the study were 6-8 week-old females except for the mice used for scRNAseq at the early timepoint that were 14-week-old females.

### Plasmid construction

H-RasV12-shp53-(IRES)-GFP-(2a)-Firefly-luciferase vector was generated from the H-RasV12-shp53-(IRES)-GFP^37^ construct using the Gibson Assembly technique^78^. The terminal IRES-GFP region of the initial vector and a P2A-luciferase2 sequence were both flanked with a shared sequence overlap and amplified by PCR. They were then inserted into SalI and PmlI sites of the initial vector.

Four oligonucleotide sequences for the miR-NRF2-based shRNAs targeting have been designed using the Block-iT RNAi designer tools (Invitrogen; Supplementary Table 4) and cloned into the pcDNA6.2-GW /EmGFPmiR plasmid according to the manufacturer protocol (BLOCK-iT polII miR RNAi, invitrogen #K4936-00). miR-NRF2 #4 and a miR-ctl^79^ (Supplementary Table 4) have been further cloned by the Gibson Assembly technique in the H-RasV12-shp53-(IRES)-GFP-(2a)-Firefly-luciferase vector.

### Stereotaxic injection

We stereotaxically performed lentiviral intracranial injection of mice to induce *de novo* tumorigenesis. The mice were anesthetized with isoflurane (2-3%, 1 L/min oxygen), and subcutaneously injected in the head with lidocaine (60 μL, 2.133 mg/mL). Analgesia was injected intraperitoneally (i.p.) presurgery and up to 24h after surgery (buprenorphine, 100 μL, 15 μg/mL). The HRasV12-shp53-(IRES)-GFP (lv) or the HRasV12-shp53-(IRES)-GFP-(2a)- Firefly-luciferase lentivirus (lv-luc, 1 μL, 6 x 10^8^ PFU/mL) was injected in the right subventricular zone (SVZ) of the brain (x=1mm, y=1mm, z=-2.3mm from the bregma). We used a Hamilton 30G needle with a silica fiber tip (MTI-FS) and an automatic injector (Harvard Apparatus). After injection, the skin wound was closed with surgical glue (SurgiBond®) and animals were placed under an infrared lamp until they recover a vigil state. From the next day, mice were injected i.p. with tamoxifen (TMX, 20 mg/mL in corn oil, Sigma #T5648-1Gi and Sigma #C8267) once per day for five consecutive days to induce the recombination of the *Pten* locus and of the *loxP-RFP-loxP* cassette of the lentivirus allowing the expression of *H-RasV12*.

### Bioluminescence imaging

We monitored tumor growth by *in vivo* bioluminescence twice a week from 14 days post intracranial injection. The mice were i.p. injected with Xenolight D-Luciferin (100 μL, 30 mg/mL, Perkin Elmer #122799), anesthetized with isoflurane and their head and back were shaved. Bioluminescence was recorded with an IVIS Spectrum In Vivo Imaging System (Perkin Elmer) and ratio were measured by normalizing the head signal on the back signal. Onset of tumor growth corresponds to a head/trunk bioluminescence ratio of 2 (see below) for the p16-3MR+vhc and p16-3MR+GCV mice and to a head bioluminescence signal of 3e10^6^ for miR-ctl and miR-NRF2 GBM-bearing mice. The difference in the evaluation of tumor growth was due to a point mutation in the P2A sequence in the HRasV12-shp53-(IRES)-GFP-(2a)-Firefly-luciferase vector.

### Mouse treatments

Mice were treated with vehicle (PBS, DMSO 20%, Sigma #D8418-50ML) or GCV (25 mg/kg/day, Selleckchem, #S1878) prepared in PBS, 20% DMSO at 21 days post injection (DPI). During the course of the study, we implemented bioluminescence-monitored GBM growth for the two paradigms p16-3MR+vhc vs p16-3MR+GCV and the WT+vhc vs WT+ABT263 (see below). The mice were treated when head to back bioluminescence ratio was superior or equal to 2 (around 24 DPI; n=43). GCV was administered via daily i.p. injections for 5 consecutive days per cycle, for two cycles with a 2-week interval between the two cycles. ABT263 (Selleckchem, #S1001) was prepared as previously described^13^ and was administered to mice by gavage at 50 mg/kg/day for 5 days.

### Kaplan-Meier mice survival studies

Kaplan-Meier survival analysis was done using Prism 8 (Graphpad software). In accordance with EU guidelines, mice were sacrificed when reaching end points (20% body weight decrease, deterioration of general condition). Mice were injected by batch. One batch always included control and experimental mice injected the same day. When control mice survival extended more than 50-52 DPI, the entire batch was removed from the analysis to exclude technical bias linked to intracranial injection.

### Mice brain collection

When reaching end points, mice were sedated with CO_2_ inhalation followed by an intracardiac perfusion with cold HBSS 1X. After harvesting the brain, GFP+ tumor was cut into two parts, under MZFL II stereomicroscope (Leica). Anterior part of the GFP+ tumor and the GFP-parenchyma were chopped and stored in TRI-reagent (Molecular Center Research, #TR 118) at −80°C or directly snap-frozen in liquid nitrogen for RNA isolation. Posterior part was snap-frozen in dry ice cooled-isopentane for histological studies. Brains were cryosectioned at 12-μm thickness (Leica cryostat).

### SA-β-gal and immunohistochemical staining

For SA-β-gal staining, brain or GBM sections were fixed in 2% PFA, 0.02% glutaraldehyde (Sigma, #340855) 10 min at RT, washed twice in PBS pH 7.0 and once in PBS pH 5.5 for 30 min. Slides were incubated in the X-gal solution as previously described^80^ for 5h30 at 37°C for mouse sections and overnight (O/N) for patient GBM sections. Slides were then washed in PBS and post-fixed in 4% PFA 10 min at RT.

For Immunohistochemical staining, brain or GBM sections were fixed in 4% PFA 10 min and washed in PBS. Endogenous peroxidases were inactivated in 1% H_2_O_2_ (in H_2_O) solution for 5 min and sections were incubated in the blocking solution (PBS 1X, 10% NGS, 3% BSA and 0.25% to 0.5% Triton) for 30 min. Sections were then incubated with the primary antibody (Supplementary Table 4) in the blocking solution for either 2h at RT or O/N at 4°C. Slides were rinsed and incubated with biotinylated secondary antibodies for 45 min at RT. An amplification step was performed using VECTASTAIN Elite ABC HRP Kit (Vector Laboratories, #PK-6100-NB) for 30 min at RT and staining was revealed by a DAB reaction. Images were acquired using an Axio Scan.Z1 (Zeiss) and extracted using the ZEN 2.0 blue edition (Zeiss) software.

### Surface area quantification

Quantifications were performed using Fiji software^81^. Region of interest (ROI) corresponding to the tumor, was selected using the ellipse tool. IHC images were then color deconvoluted according to the “Giemsa” or “H DAB” vector to assess a threshold of the SA-β-gal or DAB signal respectively. The signal threshold was adjusted in order to remove the unspecific background signal without clearing the specific one. Number of pixels was measured and the values were normalized on the GFP+ tumor surface area. For mice brain tissue, four slides with three sections on each (n=12) for SA-β-gal quantification and three slides with three sections on each (n=9) for IHC quantification were analyzed per sample. For patient sample, 4 sections were analyzed for SA-β-gal quantification.

### Western-Blotting

Total proteins were extracted from tumor samples following TRI-reagent protocol (Molecular Center Research, #TR 118). Protein pellets were solubilized in 1% SDS, 10M urea and stored at −80°C. Protein concentration was assayed using the Pierce BCA protein assay kit (ThermoFisher, #23225). Proteins were separated on 4-20% stain-free polyacrylamide gels (Mini-PROTEAN TGX Protein Gels, Bio Rad, #4568096) and transferred on a nitrocellulose membrane 0.45 μm (ThermoFisher, #88018). Membranes were probed with primary antibodies (Table S4) diluted in Super Block Blocking buffer in TBS (ThermoFisher, #37535) and incubated O/N at 4°C under gentle agitation. The secondary antibodies were incubated 1h at RT. Fluorescence was detected using the Odyssey CLx (Li-cor), specific bands were quantified using Fiji software^81^ and normalized against the corresponding β-TUBULIN band.

### Cell culture

Glioma 261 murine cell lines (GL261) were cultured in DMEM (Thermofisher #31966021) 1% foetal bovine serum (Thermofisher #A3160801). These cells were transfected with the pcDNA6.2-GW /EmGFPmiR plasmids containing miR sequences (miR-NRF2 #1, #2, #3, #4 and miR-ctl ^79^ using the FUGENE HD transfection reagent (Promega #E2311). Seven days later GFP positive cells were isolated by flow cytometry (Biorad S3e cell sorter), cultured for two more days and their total RNAs were extracted.

### RNA extraction and RT-qPCR

Total RNAs were extracted from tumor and parenchyma samples and GL261 cells following either the TRI-reagent (Molecular Center Research, #TR 118), the Maxwell RSC simplyRNA Tissue (Promega, #AS1340) and the Macherey-Nagel Mini kit Nucleospin protocol (Macherey-Nagel, #740955.50).

cDNAs were generated using the Maxima 1str cDNA Synth Kit (LifeTechnologies, K1642). Quantitative PCR was performed using LightCycler 480 SYBR Green I Master Mix (Roche, #4707516001) on a LightCycler® 480 Instrument II (Roche). Samples were run in duplicate or triplicate, transcript levels were normalized to TBP and GAPDH and analysis was performed using the 2^-ΔΔCT^ method^82^. Primers used in this study are listed in Supplementary Table 4.

### Tumor dissociation for scRNAseq

After brain harvest, GFP+ tumors were dissected under a Leica MZFL II stereomicroscope. Tumor pieces were chopped and incubated 5 min at 37°C in a HBSS-papain based lysis buffer (Worthington PAP) containing DNAse (0.01%, Worthington #LS002139) and L-Cystein (124 μg/mL, Sigma #C78805). Papain digestion was inhibited by ovomucoid (70 μg/mL, Worthington #LS003085). Tissue was further dissociated mechanically and centrifuged 300 g, 10 min at 4°C. Cells were resuspended in cold HBSS, a debris removal step was performed (Miltenyi #130-109-398) and blood cells were removed using a blood lysis buffer (Roche 11814 389001).

### Bulk RNA-seq and analysis

The quantity and quality of the total RNAs extracted were assessed by the Tapestation 2200 (Agilent), and sequenced with the Illumina NextSeq 500 Sequencing system using NextSeq 500/550 High Output Kit v2 (150 cycles, # 20024907), 400 millions of reads, 50Gbases.

Quality of raw data was evaluated with FastQC. Poor quality sequences were trimmed or removed with Fastp software to retain only good quality paired reads. Star v2.5.3a was used to align reads on mm10 reference genome using default parameters except for the maximum number of multiple alignments allowed for a read which was set to 1. Quantification of gene and isoform abundances were done with rsem 1.2.28 on RefSeq catalogue, prior to normalisation with edgeR bioconductor package. Finally, differential analysis was conducted with the glm framework likelihood ratio test from edgeR. For malignant samples, a batch effect was detected in PCA representation. To correct it, we performed the analysis by using an additive model which includes this batch variable. Multiple hypothesis adjusted p-values were calculated with the Benjamini-Hochberg procedure to control FDR.

Functional enrichment analysis was performed with clusterProfiler (v3.14.3) bioconductor package on the differentially deregulated genes with over-representation analysis (enricher function) and on all the genes with Gene Set Enrichment analysis (GSEA function; ^83^). Hallmark, Transcription factor targets (TFT) and Canonical pathways (CP) gene sets from MSigDB collections have been used, completed with some custom gene sets (Supplementary Table 1).

### Single-cell RNA-seq and analysis – 10X data

Cells suspension of four dissociated GBMs (2 WT+GCV and 2 p16-3MR+GCV) were loaded with the Chromium Next GEM Chip G Single Cell Kit (10X Genomics, #PN-1000120) and a library was generated using Chromium Next GEM Single Cell 3’ Reagent Kits v3.1 (10X Genomics, #20012850). The library was sequenced on an Illumina NovaSeq 6000 instrument using a 100 cycle S2 flow cell in XP mode, with the following parameters: 2050 million reads depth, 200 Gbases per run and 50 000 reads per cell.

The Cell Ranger Single-cell Software suite (3.0.2) was used to process the data. First, a custom reference genome was created with the mkref function to include 3’LTR and 3MR sequences into the mm10 reference genome. Count function was used on each GEM well that was demultiplexed by *mkfastq* to generate gene-cell matrices. Then, filtered_feature_bc_matrix output was loaded into Seurat bioconductor package v3.2.3 to filter the datasets and identify cell types using R v3.6. Genes expressed in at least 5 cells and cells with at least 200 features were retained for further analysis. To remove likely dead or multiplet cells from downstream analyses, cells were discarded when they had less than 500 UMIs (Unique Molecular Identifiers), greater than 60 000 UMIs, or expressed over 8 % mitochondrial genes.

All samples were merged together for downstream analysis. As no batch effects were observed among the four samples, no integration step was performed. Gene expression matrix was normalized using the negative binomial regression method implemented in the Seurat *SCTransform* function, via selection of the top 3000 variable genes and regressed out the mitochondrion expression percentage. The final dataset was composed of 20 293 genes and 26 237 cells.

To cluster cells, we computed a Principal Components Analysis (PCA) on scaled variable genes, as determined above, using Seurat’s *RunPCA* function, and visualized it by computing a Uniform Manifold Approximation and Projection (UMAP) using Seurat’s *RunUMAP* function on the top 30 PCs. We also computed the k-nearest neighbor graph on the top 30 PCs, using Seurat’s *FindNeighbors* function with default parameters, and in turn used Seurat’s *FindClusters* function with varying resolution values. We chose a final value of 0.5 for the resolution parameter at this stage of clustering. Clusters were assigned preliminary identities based on expression of combinations of known marker genes for major cell types. TME clusters were identified with the expression of *Ptprc* (*Cd45*) gene marker. In order to better identify other cell types, TME cells were removed and a second clustering with a resolution 0.6 was applied.

The *FindMarkers* function with the default parameters (min.LogFC = 0.25, min.pct = 0.25, test.use = Wilcox) was used to identify differentially expressed genes in different conditions : (i) p16-3MR+GCV vs WT+GCV in each cluster; (ii) cells from astrocyte and NP clusters with *Cdkn2a* expression ≥ 4 (307 cells) vs all the other cells (10 280 cells) in WT+GCV GBMs. Functional enrichment analysis were done with clusterProfiler (v3.14.3) bioconductor package on the differentially deregulated genes with over-representation analysis (enricher function) and with Gene Set Enrichment analysis (GSEA function^83^) on all the genes. Copy number variations (CNVs) were inferred with inferCNV package (v. 1.6.0) with the following parameters: “denoise” and a value of 0.1 for “cutoff”.

### Signature expression analyses

We analyzed the senescence signature through our tumoral SCT normalized dataset and three datasets corresponding to patient GBMs^23,26,53^. This first dataset was retrieved via the single cell portal (singlecell.broadinstitute.org) and processed via Seurat (v3.2.3), 10X samples were normalized via SCT method and Smartseq2 samples were retrieved in log2(TPM+1). Finally, we retrieved the normalized expression matrix of Johnson and colleagues via synapse (https://www.synapse.org/#!Synapse:syn22257780/wiki/604645). For both murine and patient GBM datasets, we filtered out transcriptomes expressing the *CD45* (*PTPRC*) gene and pediatric GBMs^23^ and we calculated a senescence score resulting from the single-sample GSEA^84^ using the R package GSVA version 1.40.1. For these two datasets, we computed the z-scores of the resulting enrichment scores and sliced the signature score distribution into deciles to determine the HIGH senescence cells (last decile), the LOW senescence cells (1st decile), and the others with an average senescence potential (MEDIUM).

### Cox regression analysis

Normalized intensities from TCGA microarray data were obtained from cBioPortal (cbioportal.org), filtering for GBM TCGA, Firehose Legacy dataset. First, single-sample GSEA score were calculated with gsva R package v1.32.0 for senescence genes signature and NRF2 targets signature. Secondly, we fitted a Cox proportional hazards regression model with the *coxph* function from survival R package (v2.44-1.1), with additional covariates such as p16 copy number alteration (CNA) status, age of patients. Plots were done with *ggforest* R function.

### Statistical Analysis

Data are presented as mean with standard error to the mean (SD) unless otherwise specified. Statistical comparisons were performed using Wilcoxon-Mann-Whitney, p-values unless otherwise specified (*, p<0.05; **, p<0.01; ***, p<0.001). For Kaplan-Meier survival curves, statistical significance was determined by Mantel-Cox log-rank test (*, p<0.05).

## Data availability

Generated data in this study are available through the Gene Expression Omnibus (GEO: GSE168014, GSE168040).

## Acknowledgements

The authors acknowledge the service of the ICM platforms: iGenSeq, iVector especially Phillippe Ravassard, icm-QUAN-Imaging, Pheno-ICMice, Histomics-Histology, Celis-Cell culture, the service of the animal facility UMS28 (Sorbonne University) and the service of the flow cytometry platform CYTO-ICAN especially Florence Deknuydt (Hôpital Pitié-Salpétrière). The authors are grateful to Judith Campisi, Inder Verma, Magdalena Götz, Anton Berns and the Centro Nacional de Investigaciones Oncológicas (CNIO) for providing reagents and Carol Schuurmans for comments on the manuscript. We also thank members of the Huillard-Sanson laboratory for discussions, and help with the experiments, in particular Sophie Paris for teaching the intracranial injection. This work was supported by institutional fundings from Paris Brain Institute (ICM), the Institut National de la Santé Et de la Recherche Médicale and the Centre National de la Recherche Scientifique and grants from the ATIP-AVENIR (EH), the Cancéropole Ile de France (Emergence Program, ILR), the Ligue contre le cancer, comité Ile de France (ILR); the Fondation ARC pour la Recherche sur le Cancer (EH, ILR); the SIRIC-CURAMUS (joint Emergence program ILR and CA). R. Salam was supported by fellowships from the French Ministry of Education and Research and the Ligue Nationale Contre le Cancer; A. Saliou was supported by a fellowship from the Ligue Nationale Contre le Cancer.

## Author contributions

RS: Conceptualization, Methodology, Experiments, Formal analysis, Methodology, Writing - Original Draft. AS: Conceptualization, Methodology, Experiments, Formal analysis, Methodology, Writing - Original Draft. FB: Methodology, Formal analysis, Resources. MB: Formal analysis, Bioinformatics. CA: Formal analysis, Bioinformatics, Funding acquisition, Writing - Original Draft. CC: Resources. AA: Bioinformatics. LC: Resources. MS: Funding acquisition, Resources. EH: Funding acquisition, Resources. LB: Formal analysis, Bioinformatics, Writing - Original Draft. JG: Formal analysis, Bioinformatics, Writing - Original Draft. ILR: Conceptualization, Methodology, Experiments, Formal analysis, Methodology, Funding acquisition, Supervision,

## Materials & Correspondence

Further information and material requests should be addressed to Isabelle Le Roux.

