## Supplementary material for "Cellular senescence in malignant cells promotes tumor progression in mouse and patient Glioblastoma": Supplen

**Supplementary data**

### Supplementary Figure 1

**a**

| #Tumor | Type | Molecular alterations/genes | Molecular alterations/chromosomes | p16 status | %SA-β-gal area |
|---|---|---|---|---|---|
| 1 | GBM | hTERT C228T | 7 gain; 9p loss; 10 loss | deleted | 1.5 |
| 2 | GBM | EGFR ampl.; hTERT C228T; PTEN mut.; EGFR mut. | 7 gain; 9p partial loss; 10 loss; 19 gain | WT/normal | 3.17 |
| 3 | GBM | no alteration | no alteration | WT/normal | 0.57 |
| 4 | GBM | hTERT C250T; MDM2 ampl. | no alteration | WT/normal | 0.27 |
| 5 | GBM | hTERT C228T; TP53 mut.; RB1 mut.; CDK4 ampl. | 7 gain; 10 loss | WT/normal | 1.95 |
| 6 | GBM | hTERT C228T; TP53 mut.; PTEN mut.; PDGFRa ampl. | 7 gain; 9p partial loss; 10 loss | deleted | 0.06 |
| 7 | GBM | hTERT C228T; TP53 mut.; EGFR ampl.; MGMT methyl. | 7 gain; 9p loss; 10 loss | deleted | 0.34 |
| 8 | GBM | hTERT C228T; TP53 mut.; PTEN mut.; ATRX mut.; MDM2 ampl.; MGMT ampl.; CDK4 ampl. | 7 gain; 10 loss | WT/normal | 0.59 |
| 9 | GBM | TP54 mut.; MGMT methyl. | 1p partial del.; 7 gain; 9p loss; 10 loss | deleted | 1.32 |
| 10 | GBM | EGFR ampl.; hTERT C250T; TP53 mut. | 7 gain; 9p partial loss; 10 loss; 9p19q gain | deleted | 0.15 |
| 11 | GBM | hTERT C250T; MGMT methyl.; FUBP1 mut.; ATRX mut.; NF1 mut. | 9p loss; 10 loss | heterozygous | 3.25 |
| 12 | GBM | MGMT methyl. | 7 gain; 4q partial loss; 10 loss; 13q partial loss; 19 gain | WT/normal | 1.27 |
| 13 | GBM | no alteration | 7 gain; 9p partial loss; 10 loss; 14q loss; 15q partial loss; 17q partial loss | WT/normal | 0.04 |
| 14 | GBM | EGFR ampl.; hTERT C228T; MGMT methyl. | 1 partial loss; 7 gain; 9p partial loss; 10 loss; 19 gain | deleted | 0.05 |
| 15 | AA IV | IDH1 mut.; TP53 mut.; ATRX mut.; NF1 mut.; PTEN mut.; PMS2 mut.; CDKN2A H83Y | 7 gain; 9p partial loss; 10q partial loss | mutated | 1.55 |
| 16 | AA III | IDH1 mut. | NS | NS | 0.15 |
| 17 | AA III | IDH1 mut.; TP53 mut. | 1p gain; 7q gain; 19q gain | WT/normal | 0.56 |
| 18 | A II | IDH2 mut.; TP53 mut.; ATRX mut. | no alteration | WT/normal | 0.85 |
| 19 | AA III | IDH1 mut.; TP53 mut.; ATRX mut. | 1p partial del. | WT/normal | 2.62 |
| 20 | O III | IDH1 mut.; hTERT C250T; CIC mut. | 1p19q codel.; 9p loss | heterozygous | 1.97 |
| 21 | O III | IDH1 mut.; hTERT C250T; CIC mut. | 1p19q codel.; 8 loss | WT/normal | 0.36 |
| 22 | O III | IDH1 mut.; hTERT C228T; CIC mut. | 1p19q codel. | WT/normal | 0.78 |
| 23 | O III | IDH1 mut.; hTERT C228T; PIK3CA mut. | 1p19q codel.; 6p partial loss; 9p partial loss; 15q loss; 18 loss; 22q partial loss | deleted | 0.05 |
| 24 | O III | IDH1 mut.; hTERT C228T; PTEN mut. | 1p19q codel. | WT/normal | 6.77 |
| 25 | O III | IDH1 mut.; hTERT C228T | 1p19q codel.; 14q partial loss | WT/normal | 0.04 |
| 26 | O III | IDH2 mut.; hTERT C250T; CIC mut. | 1p19q codel. | WT/normal | 0.26 |
| 27 | O II | IDH1 mut.; CIC mut. | 1p19q codel. | WT/normal | 0.38 |
| 28 | O II | IDH2 mut.; hTERT C228T | 1p19q codel. | WT/normal | 0.48 |

recurrence

senescent category
>1%
0.1-1%
≤0.1%

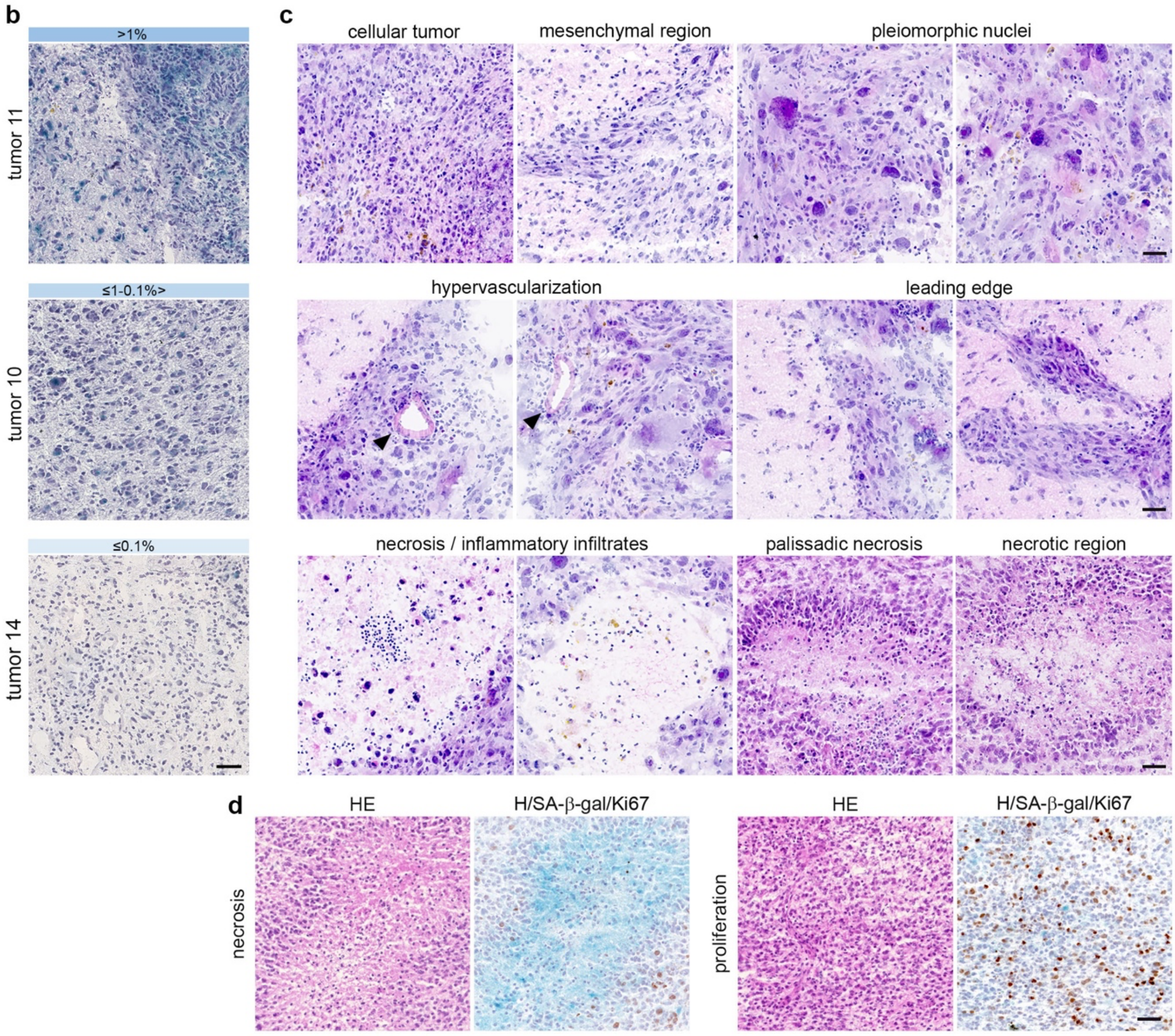

**Supplementary Figure 1. Identification of senescent cells in patient and mouse gliomas.**
(**a**) Table recapitulating the patient resected diffuse gliomas stained for SA-β-gal and annotated for their glioma type, molecular alterations and percentage of SA-β-gal area over the tumor area. Recurrent gliomas are highlighted in grey.
(**b**) Representative SA-β-gal staining (blue) of the 3 categories of senescence in patient gliomas. Sections are counterstained with hematoxylin (H).
(**c**) Representative Hematoxylin and Eosin (HE) staining on mouse GBM cryosections. The mouse GBM model recapitulates the patient GBM histological features.
(**d**) Representative HE and SA-β-gal/Ki67 staining on adjacent mouse GBM cryosections highlighting the presence SA-β-gal+ cells in proliferative and necrotic areas.
Scale bars, **b**: 50 μm; **c**: 20 μm; **d**: 100 μm. amp: amplification; del.: deletion; methyl.: methylation; mut.: mutation; NS: not specified.

**Supplementary Figure 2**

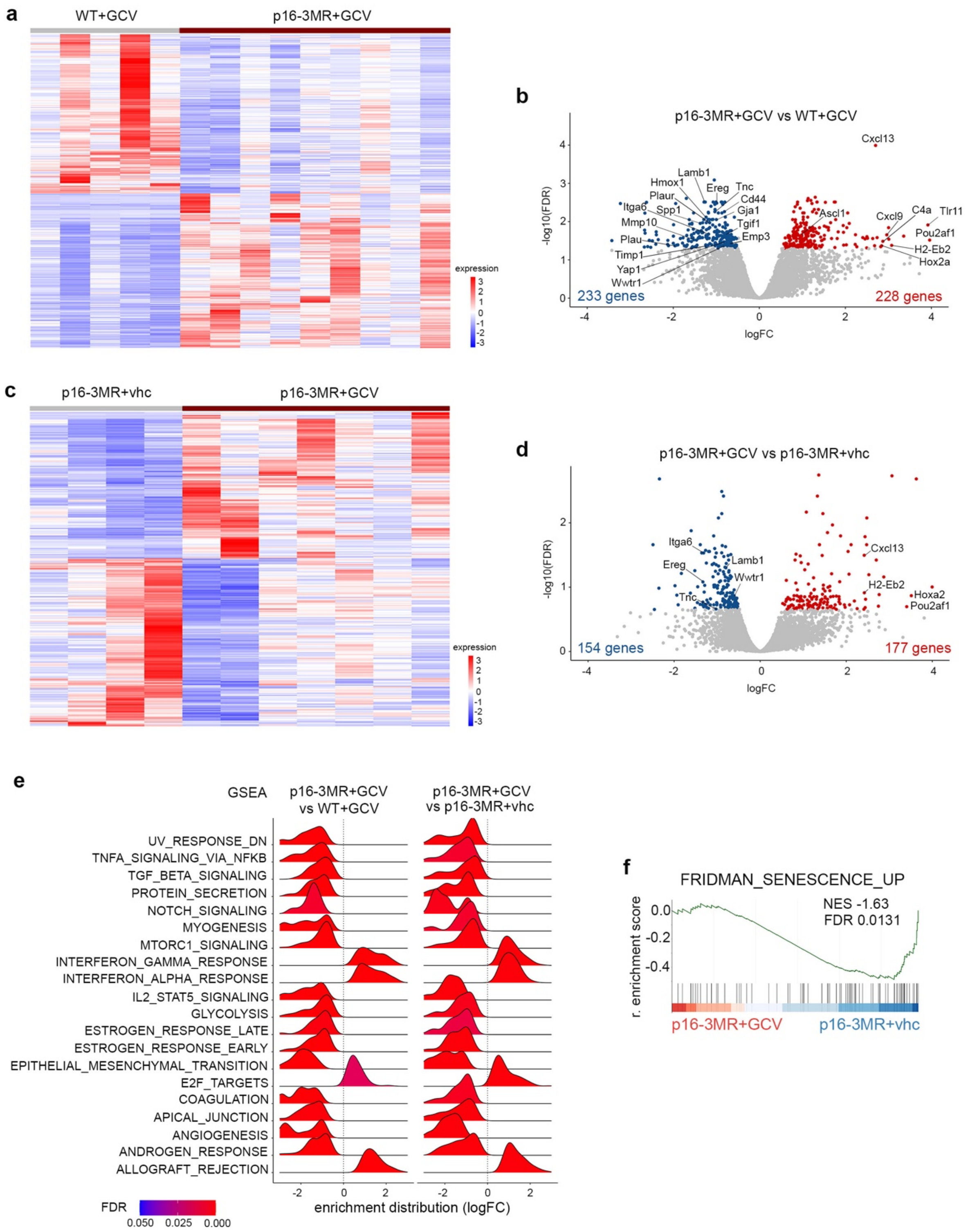

**Supplementary Figure 2. Senescent cells partial removal increases the survival of GBM bearing mice**

(**a**) Heatmaps representing bulk RNAseq analysis of the differentially expressed (DE) genes (FDR<0.05; logFC>0.5) between p16-3MR+GCV (n=9) compared with WT+GCV (n=5) GBMs collected at the end points of the mice.

(**b**) Volcano plots of the DE genes (FDR<0.05; logFC>0.5) between p16-3MR+GCV (n=9) compared with WT+GCV (n=5) GBMs collected at the end points of the mice.

(**c**) Heatmaps representing bulk RNAseq analysis of the DE genes (FDR<0.05; logFC>0.5) between of p16-3MR+GCV (n=7) compared with p16-3MR+vhc GBMs (n=4) collected at the end points of the mice.

(**d**) Volcano plots of the DE genes between p16-3MR+GCV (n=7) compared with p16-3MR+vhc GBMs (n=4) collected at the end points of the mice. Annotated genes are common to those in B.

(**e**) GSEA ridge plots of the significant representative Hallmark gene lists common to the two RNAseq analysis (**a**, **b** and **c**, **d**).

(**f**) GSEA graph representing the enrichment score of the Fridman senescence pathway in p16-3MR+GCV compared with p16-3MR+vhc GBMs.

GCV: ganciclovir; FDR: false discovery rate; NES: normalized enrichment score; r. enrichment score: running enrichment score.

### Supplementary Figure 3

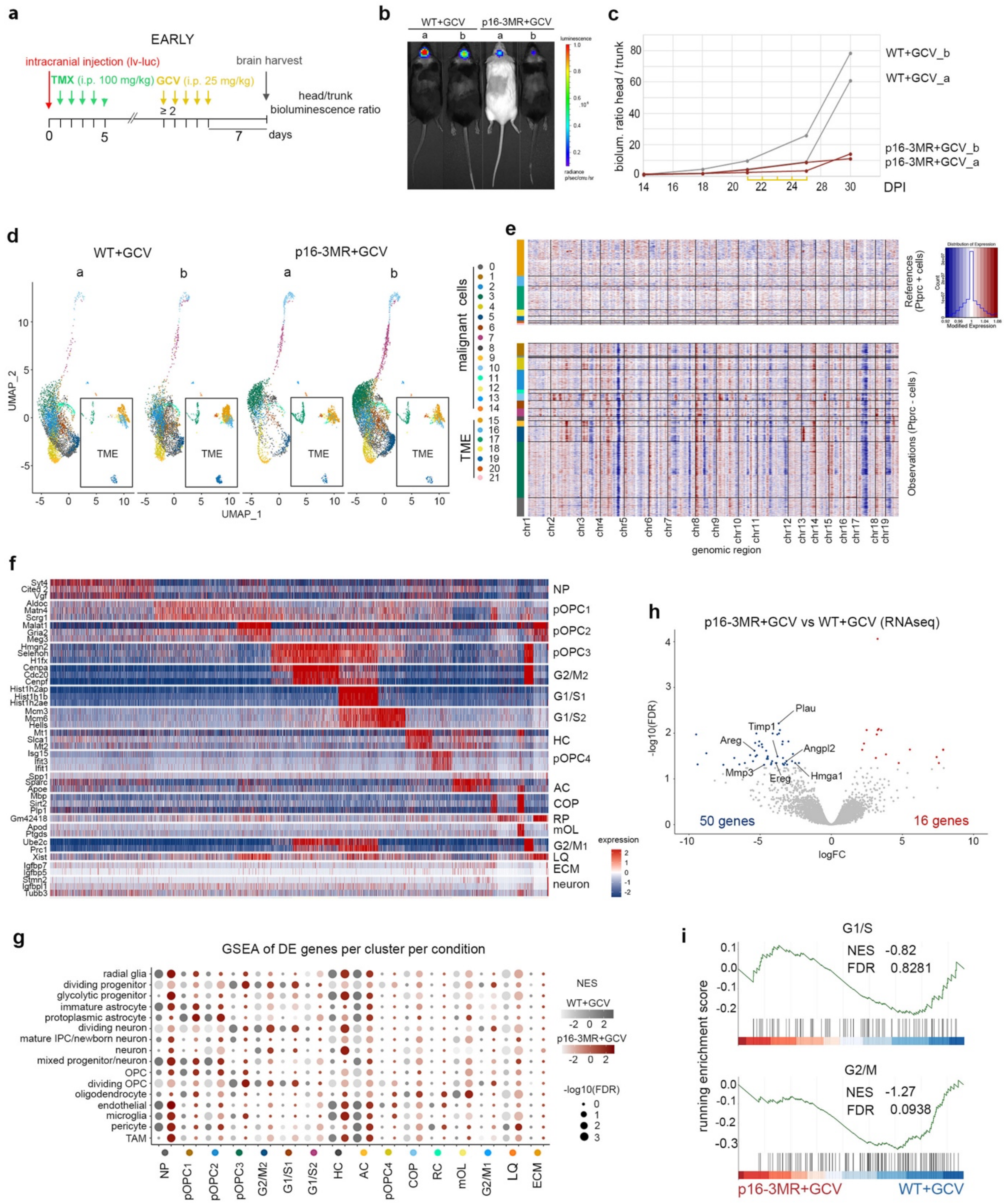

**Supplementary Figure 3. Identification of p16$^{Ink4a\ Hi}$ cells in a subset of malignant cells**

(**a**) Timeline of the mouse GBMs generation for scRNAseq at the early timepoint.

(**b**) *In vivo* bioluminescence imaging of WT+GCV and p16-3MR+GCV GBM-bearing mice.

(**c**) Graph representing the head to body ratio bioluminescence of GBM-bearing mice over time. Yellow lines correspond to GCV injections.

(**d**) UMAP plots of WT+GCV and p16-3MR+GCV GBM cells per biological sample at a 0.5 resolution and annotated malignant cells and cells from the tumor microenvironment (TME).

(**e**) Heatmap representing inference of chromosomal copy number variations (CNVs) in WT+GCV GBMs with cells as rows, grouped in clusters, and genes as columns, ordered according to chromosome.

(**f**) Heatmap of the top 3 differentially expressed (DE) genes (FDR<0.05; avlogFC>0.25) in malignant clusters (0.6 resolution) in WT+GCV GBMs. When the same gene is DE in more than one cluster, it appears only once.

(**g**) GSEA dot plots of DE genes (FDR<0.05; avlogFC>0.25) in WT+GCV (grey dots) and p16-3MR+GCV (red dots) GBMs of gene lists from Bhaduri *et al.*[1] (Supplementary Table 1).

(**h**) Volcano plots of the DE genes between of p16-3MR+GCV (n=4) compared with WT+GCV GBMs (n=4).

(**i**) GSEA graph representing the enrichment score of the cycling pathways (Weng *et al.*[2], 2019; Supplementary Table 1) in p16-3MR+GCV (n=4) compared with WT+GCV (n=4).

**d-g:** analysis performed from scRNAseq data as shown in **a-c**. **h** and **i**: analysis performed from bulk RNAseq data at the early time point of the mice. GCV: ganciclovir; TMX: tamoxifen; i.p.: intraperitoneal; lv-luc: lentivirus-luciferase; UMAP: uniform manifold approximation and projection; chr: chromosome; FDR: false discovery rate; NES: normalized enrichment score.

### Supplementary Figure 4

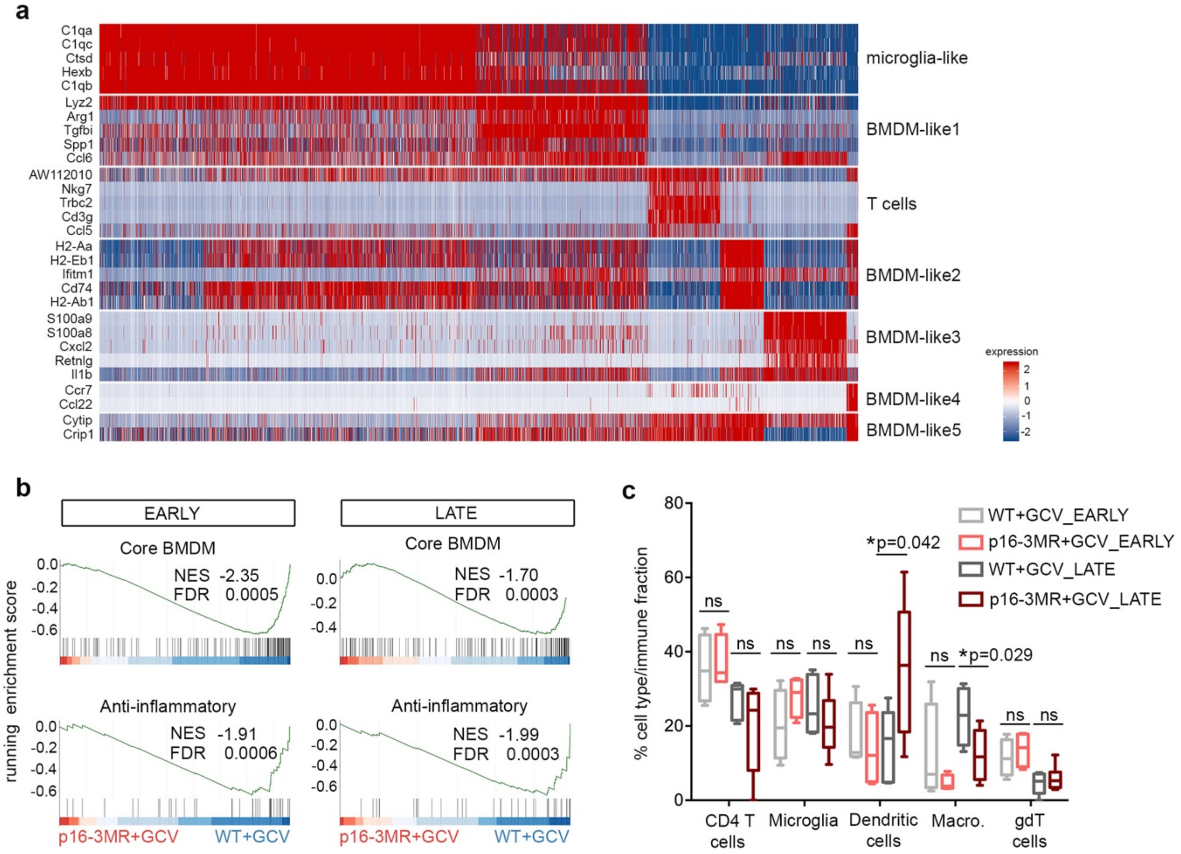

**Supplementary Figure 4. Modulation of the immune compartment following $p16^{Ink4a\ Hi}$ cells partial removal**

(**a**) Heatmap of the top 5 differentially expressed (DE) genes (FDR<0.05; avlogFC>0.25) in *CD45*+ clusters (0.5 resolution) in WT+GCV GBMs. When the same gene is DE in more than one cluster, it appears only once.

(**b**) GSEA graphs representing the enrichment score of core bone marrow-derived macrophages (BMDM) and Anti-inflammatory pathways (Bowman *et al.*[3]; Darmanis *et al.*[4]; Supplementary Table 1) in p16-3MR+GCV compared with WT+GCV GBMs at the early and late timepoints. Analysis performed from bulk RNAseq data.

(**c**) Bar plot representing the estimation of the abundance of immune cell types in WT+GCV and p16-3MR+GCV GBMs using CIBERSORT (reference data set GSE124829) at the early and late time points. Analysis performed from bulk RNAseq data. Statistical significance was determined by Wilcoxon-Mann-Whitney test (ns, not significant; *, p<0.05).

FDR: false discovery rate; NES: normalized enrichment score; macro.: macrophages; GCV: ganciclovir.

### Supplementary Figure 5

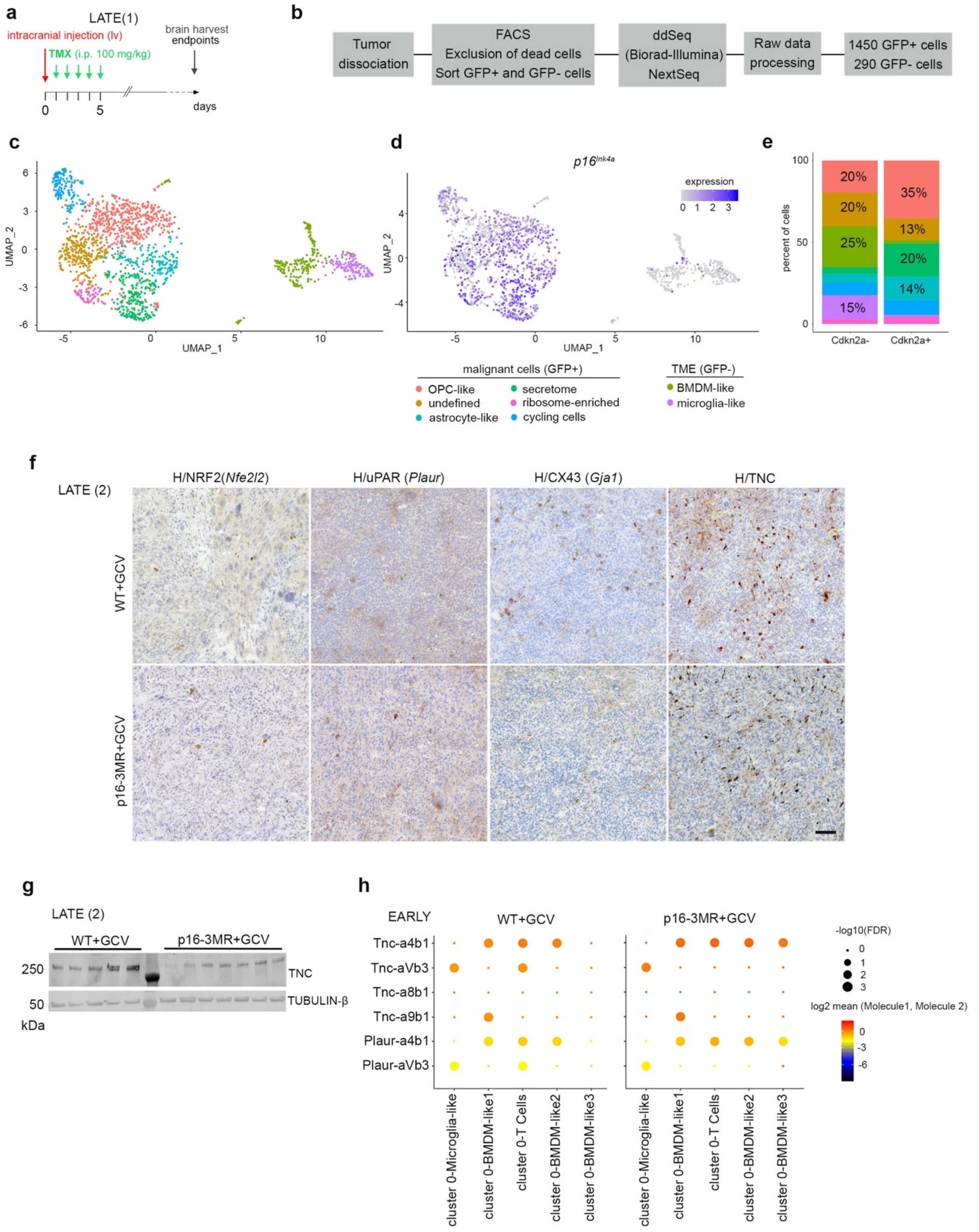

**Supplementary Figure 5. Identification of NRF2 activity and its putative targets in p16$^{Ink4a}$ $^{Hi}$ malignant cells**

(**a**) Timeline of the mouse GBM generation for scRNAseq at the late timepoint (LATE(1)).

(**b**) Scheme of the scRNAseq experiment.

(**c**) UMAP plots of GBM cells and annotated cell type at a 0.6 resolution.

(**d**) UMAP plots of the expression of *p16$^{Ink4a}$* in GBM cells.

(**e**) Barplot representing the percentage of cells per cluster positive and negative for *p16$^{Ink4a}$* expression.

(**f**) Low magnification of representative immunohistochemistry (IHC, brown) counterstained with hematoxylin (H, purple) on mouse GBM cryosections at the late timepoint.

(**g**) Western blot (WB) for TNC from independent WT+GCV (n=5) and p16-3MR+GCV (n=7) GBMs collected at the late timepoint. Each lane corresponds to one GBM. TUBULIN-β was taken as a loading reference. Quantification of the WB is shown in Fig. 5I.

(**h**) Dot plot representing ligand-receptor interactions between the cluster 0 and the immune clusters in the scRNAseq data at the early timepoint using CellPhoneDB. The colors indicate the mean expression of the ligand-receptor complexes.

Scale bar, **f**: 40 µm. GCV: ganciclovir; TMX: tamoxifen; FACS: fluorescence-activated cell sorting; UMAP: uniform manifold approximation and projection; TME: tumor microenvironment.

**Supplementary Figure 6**

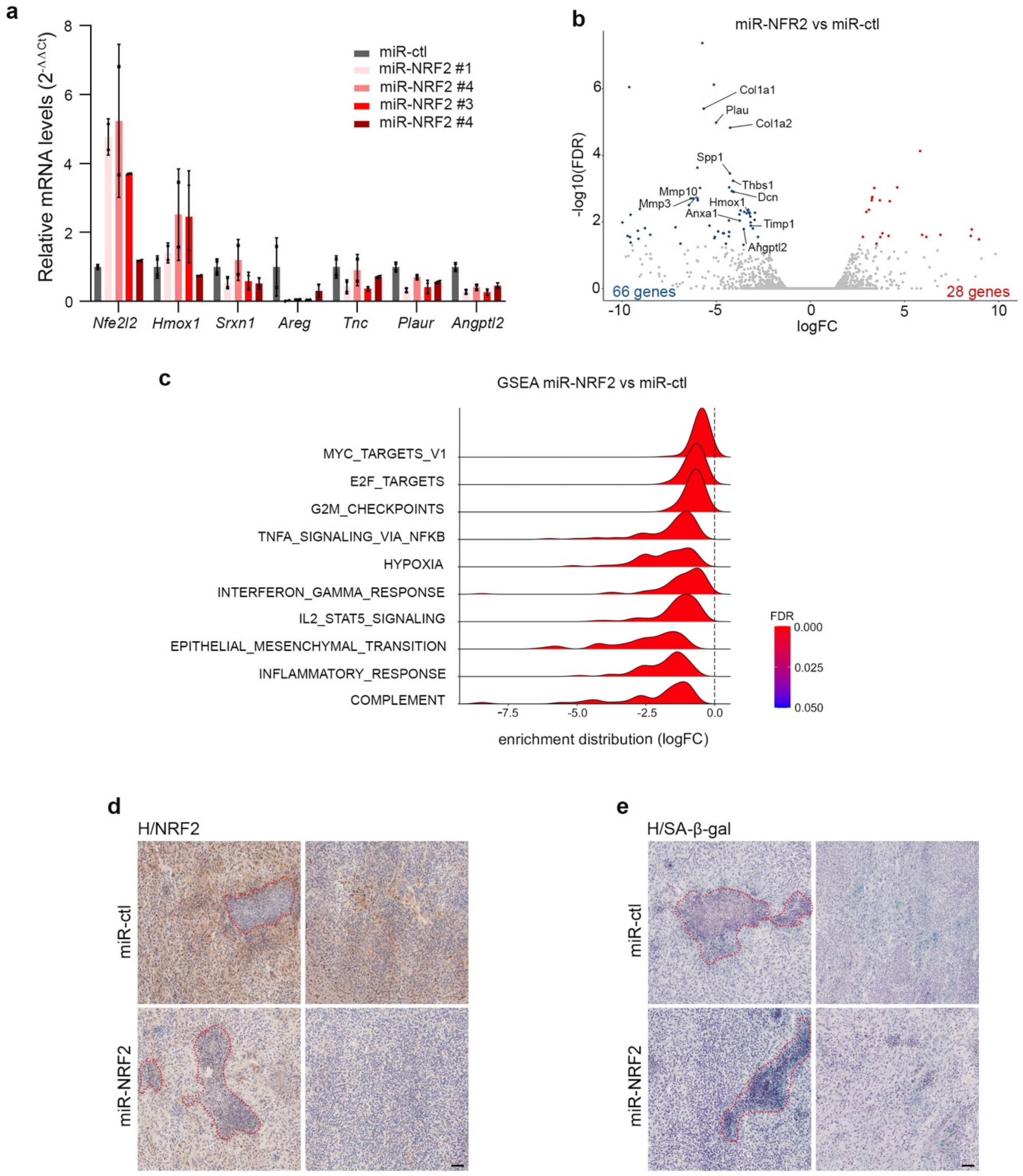

**Supplementary Figure 6. Knockdown of NRF2 in malignant cells recapitulates most features of the senolytic treatment**

(**a**) Relative transcript levels shown as ratios of normalized values of GL261 cells expressing miR-NRF2 over miR-ctl. The graph shows two independent experiments performed in duplicate. miR-NRF2 #4 reduces the expression of all the NRF2 targets (canonical targets: *Hmox1* and *Srxn1*, and NRF2 targets from the combined analysis: *Areg, Tnc, Plaur* and *Angptl2*) and was subsequently used for *in vivo* experiments.

(**b**) Volcano plots of the differentially expressed (DE) genes in miR-NRF2 GBMs (n=3) compared with miR-ctl GBMs (n=3).

(**c**) GSEA ridge plots of the 10 most significant representative Hallmark gene lists.

(**d**) Low magnification of representative NRF2 IHC (brown) counterstained with hematoxylin (purple) on miR-ctl (n=4) and miR-NRF2 (n=4) GBM cryosections.

(**e**) Low magnification of representative SA-β-gal staining (blue) counter stained with hematoxylin (purple) on miR-ctl (n=4) and miR-NRF2 (n=4) GBM cryosections..

Scale bar, **d**, **e** 100 µm. **d**, **e**, Necrotic areas are outlined in red dashed lines. **b**, **c**: Analysis performed from bulk RNAseq of GBMs collected at the late time point.

### Supplementary Figure 7

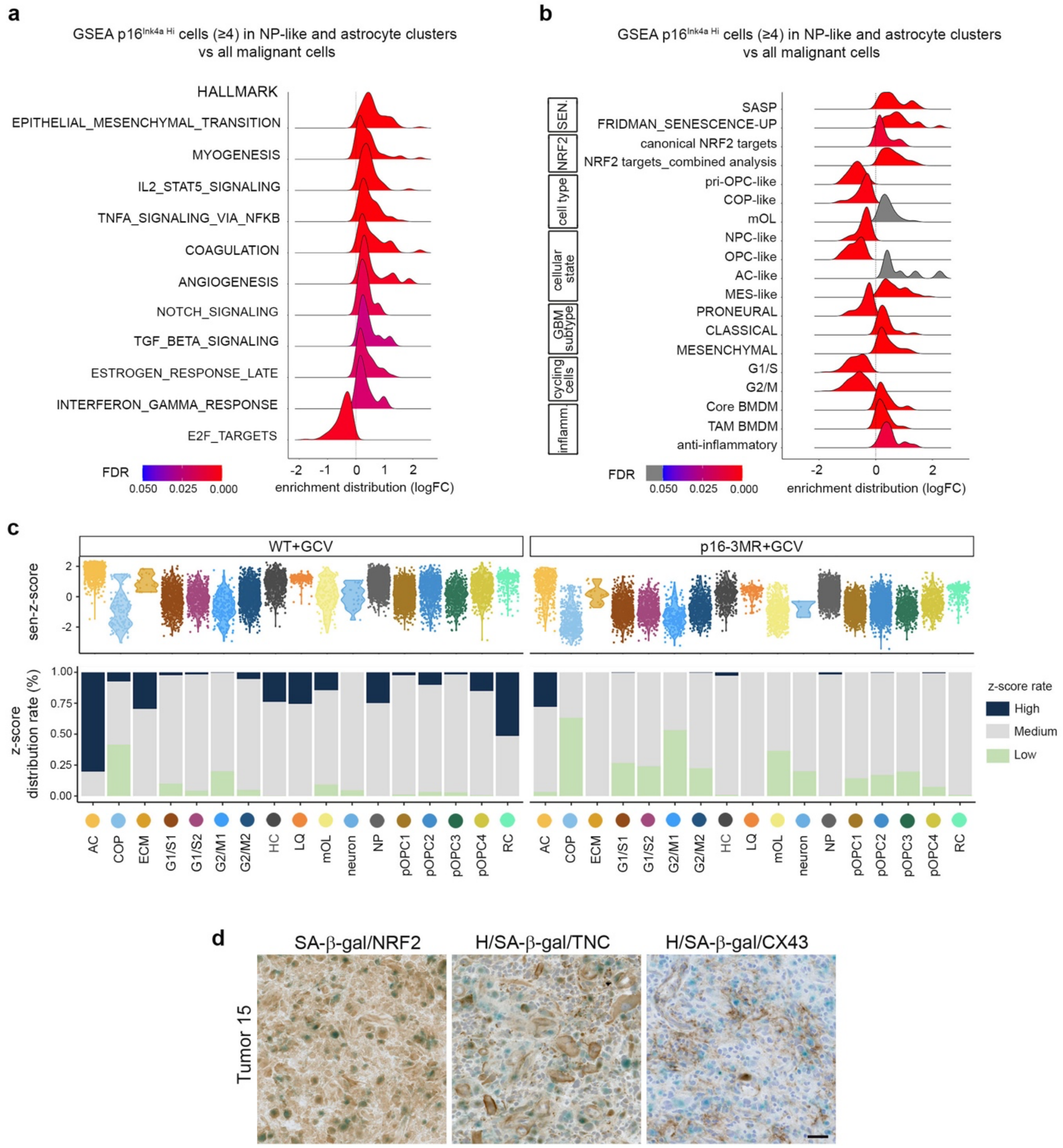

**Supplementary Figure 7. Mouse senescent signature is conserved in patient GBM and its enrichment score is predictive of a worse survival**

(**a**) GSEA ridge plot of significant Hallmark gene lists between the astrocyte and NP-like clusters compared with the remaining malignant cells in WT+GCV GBMs. The Hallmark gene lists represent the gene lists common to the GSEA analysis of the p16-3MR+GCV GBMs compared with controls (Supplementary Fig. 2e).

(**b**) GSEA ridge plot of gene lists used throughout the study (Supplementary Table 1), between the astrocyte and NP-like clusters compared with the remaining malignant cells in WT+GCV GBMs.

(**c**) Top: violin plots of the single sample GSEA (ssGSEA) senescent Z-score (sen-Z-score) in all malignant cells of WT+GCV and p16-3MR+GCV GBMs. Bottom: barplots of the percentage of the ssGSEA senescent Z-score distribution rate in all malignant cells of mouse GBMs. High and Low distribution rates correspond to the highest and lowest decile, respectively.

(**d**) Low magnification of representative SA-β-gal staining (blue) coupled with IHC (brown) and counterstained with hematoxylin (H, purple) on patient GBM cryosections. 3 patient GBMs were analyzed per antibody. Scale bar: 40 µm.

### Supplementary Methods

#### Bulk RNA-seq and analysis

CIBERSORT[5] was used to accurately quantify the relative abundances of 6 distinct immune cell types according to the ImmGen immune cell genes signature (reference GSE124829).

#### Single-cell RNA-seq and analysis – 10X data

We searched for ligand/receptor interactions between cluster 0 et *Cd45* positive clusters at 0.5 resolution in our single cell data, using CellPhoneDB v2.1.4.

#### Single-cell RNA sequencing and analysis – ddSeq data

Cell suspension of one dissociated GBM was loaded in 4 wells (3 wells with GFP+ cells or malignant cells and 1 well with GFP- cells or non-malignant cells) on the ddSEQ Single-Cell Isolator (Biorad). A library was generated using SureCell™ Whole Transcriptome Analysis 3′ Library Prep Kit for the ddSEQ System (Illumina, #20014280) and was sequenced on a Nextseq 500 Illumina sequencing system, using a High Output Kit (150 cycles), with the following parameters: 400 million reads depth, 50Gbases, and 70 million reads per sample.
Cutadapt 1.18 was used to trim nextera adapters in 3' on reads, then a quality control of sequences was done with FastQC. Cellular and UMIs barcodes were extracted with the ddSeeker (v 0.9.0) tool with default parameters. The following steps were done with Drop-seq tool (v 2.0.0). Trimming of 5' adapter sequences and of polyA tails was performed. Unaligned BAM were transformed to fastq with Picard tool, prior to alignment with STAR on mm10 reference genome. Ddseeker bam outputs previously tagged with molecular/cell barcode were merged with aligned BAM files, according to Drop-seq tool cookbook. Finally, TagReadWithGeneExonFunction was used to annotate each read with the gene it belongs to, and DigitalExpression was used to count gene transcripts in each cell. The output DGE matrix file is a matrix with a row for each gene, a column for each cell, containing the number of transcripts observed. This output was loaded into the Seurat (v2) R package for further analysis keeping only cells where at least 200 features were detected, and genes detected in at least 5 cells. The final dataset contains 1740 cells and 15 448 genes. To normalize the data, we applied the global-scaling normalization method "LogNormalize" that normalizes the feature expression measurements for each cell by the total expression, multiplies this by a scale factor (10 000 by default), and log-transforms the result. Then, highly variable genes were detected prior to scaling transformation. To cluster cells, we computed a Principal Components Analysis (PCA) on scaled variable genes, as determined above, using Seurat's *RunPCA* function, and visualized it by computing a Uniform Manifold Approximation and

Projection (UMAP) using Seurat's *RunUMAP* function on the top 10 PCs. We used *FindClusters* function with a resolution of 0.6 resulting in 8 clusters. TME and malignant clusters cells were identified according to the expression of the GFP transgene. Then, the *FindAllMarkers* function was used to extract top differentially expressed genes of each cluster, and to annotate them.
